# Structural basis for capsid recruitment and coat formation during HSV-1 nuclear egress

**DOI:** 10.1101/2020.03.11.988170

**Authors:** Elizabeth B. Draganova, Jiayan Zhang, Z. Hong Zhou, Ekaterina E. Heldwein

**Affiliations:** Department of Molecular Biology and Microbiology, Tufts University School of Medicine, Boston, MA, 02111, USA; Department of Microbiology, Immunology & Molecular Genetics, University of California, Los Angeles (UCLA), Los Angeles, California, 90095, USA; Molecular Biology Institute, UCLA, Los Angeles, California, 90095, USA; California NanoSystems Institute, UCLA, Los Angeles, California, 90095, USA

**Keywords:** HSV-1, herpesvirus, nuclear egress, nuclear egress complex, NEC, membrane budding, capsid budding

## Abstract

During herpesvirus infection, egress of nascent viral capsids from the nucleus is mediated by the viral nuclear egress complex (NEC). NEC deforms the inner nuclear membrane (INM) around the capsid by forming a hexagonal array. However, how the NEC coat interacts with the capsid and how curved coats are generated to enable budding is yet unclear. Here, by structure-guided truncations, confocal microscopy, and cryoelectron tomography, we show that binding of the capsid protein UL25 promotes the formation of NEC pentagons rather than hexagons. We hypothesize that during nuclear budding, binding of UL25 situated at the pentagonal capsid vertices to the NEC at the INM promotes formation of NEC pentagons that would anchor the NEC coat to the capsid. Incorporation of NEC pentagons at the points of contact with the vertices would also promote assembly of the curved hexagonal NEC coat around the capsid, leading to productive egress of UL25-decorated capsids.

---

To replicate, all viruses must assemble their progeny virions and release them from the cell while overcoming many obstacles, including cellular compartmentalization. Viruses are thus experts at hijacking, manipulating, and, sometimes, even remodeling cellular architecture during viral morphogenesis and egress. Identifying and understanding the unique aspects of virus-induced cellular remodeling could unveil targets for therapeutic intervention; yet, we are only beginning to understand the mechanisms behind many of these processes.

One prominent example of virus-induced remodeling of cellular architecture can be observed during egress of herpesviruses – enveloped, double-stranded DNA viruses that infect a wide range of hosts, from mollusks to humans. All herpesviruses can establish lifelong, latent infections within the host, from which they can periodically reactivate, spreading to uninfected tissues and hosts and causing a number of ailments. When the virus actively replicates during a primary infection or reactivation of a latent infection, the progeny virions are assembled and released from the cell in a process termed egress whereby herpesvirus capsids traverse cellular membranes twice [reviewed in (Johnson and Baines 2011, Bigalke and Heldwein 2016, Roller and Baines 2017)]. First, nuclear capsids bud at the inner nuclear membrane (INM) forming enveloped vesicles that pinch off into the perinuclear space. These perinuclear viral particles fuse with the outer nuclear membrane, which releases the capsids into the cytosol. Cytoplasmic capsids then bud again at vesicles derived from the *trans*-Golgi network and early endosomes [reviewed in (Johnson and Baines 2011)] to form mature, infectious virions that are released from the cell by exocytosis. Whereas many enveloped viruses acquire their lipid envelopes by budding at the cytoplasmic membranes or the plasma membrane, herpesviruses are unusual among vertebrate viruses in their ability to bud at the nuclear envelope (Bigalke and Heldwein 2016).

Capsid budding at the nuclear envelope requires two conserved herpesviral proteins, which are named UL31 and UL34 in herpes simplex virus type 1 (HSV-1), that form the nuclear egress complex (NEC) [reviewed in (Mettenleiter et al. 2013, Bigalke and Heldwein 2016, Bigalke and Heldwein 2017)]. The NEC heterodimer is anchored at the INM through the single C-terminal transmembrane helix of UL34 and faces the nucleoplasm (Shiba et al. 2000). UL31 is a nuclear phosphoprotein that colocalizes with UL34 (Chang and Roizman 1993, Reynolds et al. 2001) and interacts with the capsid during nuclear egress (Trus et al. 2007, Yang and Baines 2011). Both UL31 and UL34 are necessary for efficient nuclear egress, and in the absence of either protein, capsids accumulate in the nucleus and viral replication is reduced by several orders of magnitude (Roller et al. 2000, Fuchs et al. 2002).

Previously, we discovered that HSV-1 NEC has an intrinsic ability to deform and bud membranes by demonstrating that purified recombinant NEC vesiculates synthetic lipid bilayers *in vitro* without any additional factors or chemical energy (Bigalke et al. 2014). Similar findings were reported with the NEC homolog from a closely related pseudorabies virus (PRV) (Lorenz et al. 2015). Using cryogenic electron microscopy and tomography (cryoEM/ET), we showed that the NEC forms hexagonal “honeycomb” coats on the inner surface of budded vesicles formed *in vitro* (Bigalke et al. 2014). Very similar hexagonal coats were observed in capsidless perinuclear vesicles formed *in vivo*, in uninfected cells expressing PRV NEC (Hagen et al. 2015).

Additionally, HSV-1 NEC formed a hexagonal lattice of the same dimensions in crystals (Bigalke and Heldwein 2015). The high-resolution crystal structure of the hexagonal NEC lattice revealed interactions at the lattice interfaces (Bigalke and Heldwein 2015), and subsequent work confirmed that mutations that disrupt oligomeric interfaces reduce budding *in vitro* (Bigalke et al. 2014, Bigalke and Heldwein 2015) and *in vivo* (Roller et al. 2010, Arii et al. 2019). Collectively, these findings established the NEC as a viral budding machine that generates negative membrane curvature by oligomerizing into a hexagonal coat on the surface of the membrane.

What remains unclear, however, is how the NEC achieves appropriate coat geometry compatible with negative membrane curvature formation during budding. A purely hexagonal arrangement is flat, so curvature is typically achieved either by insertions of pentagons as found at 12 vertices of an icosahedron (Zandi et al. 2004), or by inclusion of irregular defects as observed in several viral coats (Heuser 2005, Briggs et al. 2009, Hyun et al. 2011, Schur et al. 2015). It is tempting to speculate that the capsid geometry may influence the geometry of the NEC coat. In perinuclear viral particles visualized in infected cells, the NEC coats appear to be tightly associated with the capsid (Hagen et al. 2015). Capsid interactions with the NEC during nuclear budding may be mediated by binding of the capsid protein UL25 to UL31 (Yang and Baines 2011, Yang et al. 2014). Moreover, UL25 forms pentagonal complexes at the vertices of the icosahedral herpesvirus capsids (Furlong 1978, Dai and Zhou 2018). However, it is unknown whether interaction with a mature capsid could promote pentagonal formation within NEC coats and if so, how the NEC coat would be arranged around the capsid.

A fortuitous observation that HSV-1 UL25, a capsid protein that decorates the vertices, co-localizes with synthetic liposomes *in vitro* in the presence of the NEC prompted us to investigate interactions between UL25 and NEC and the effect of UL25 on NEC-mediated budding *in vitro*. Here, by confocal microscopy, we show that free UL25 (i.e., not on capsid vertices) inhibits NEC-mediated budding *in vitro*. 3D visualization of the molecular architecture by cryoET further reveals that free UL25 forms a net of interconnected five-pointed stars on top of membrane-bound NEC layer that may block budding by preventing membrane-bound NEC coats from undergoing conformational changes required for budding. We also found that the NEC forms an alternative pentagonal, rather than hexagonal, arrangement when bound to the UL25, and that this phenomenon requires residues 45-73 that form the UL25/UL25 helical bundles on the native capsids. We hypothesize that during nuclear budding, NEC pentagons formed at the points of contact with the capsid vertices both help anchor the NEC coat to the capsid and generate appropriate coat curvature through the inclusion of pentagons into a hexagonal coat as it assembles around the capsid. This mechanism would ensure successful budding and egress of the UL25-decorated viral capsid.

## RESULTS

### Generation of UL25 variants

HSV-1 UL25 can be expressed in soluble form in *E. coli* only when residues 1-44 are deleted (Bowman et al. 2006). Residues 1-50 are necessary and sufficient for capsid binding (Cockrell et al. 2009), and in the cryoEM structure of the HSV-1 capsid, these residues mediate extensive interactions with another capsid protein, UL17 (Dai and Zhou 2018). This suggests that these residues are likely disordered in free UL25, potentially leading to aggregation and poor solubility. Therefore, we generated and expressed an HSV-1 UL25Δ44 construct, which lacks residues 1-44. UL25Δ44 was soluble and could be purified, in agreement with the previous report (Bowman et al. 2006), but was proteolytically cleaved during purification despite the presence of protease inhibitors (Fig. 1b). N-terminal sequencing (data not shown) of the cleavage product revealed that UL25Δ44 was cut between residues Q72 and A73. To prevent heterogeneity due to cleavage, we generated two constructs: UL25Δ44 Q72A, which has a single point mutation that should eliminate the cleavage site, and UL25Δ73, which corresponds to the cleavage product. Both constructs yielded a single UL25 species after purification (Fig. 1b).

**Fig. 1.**
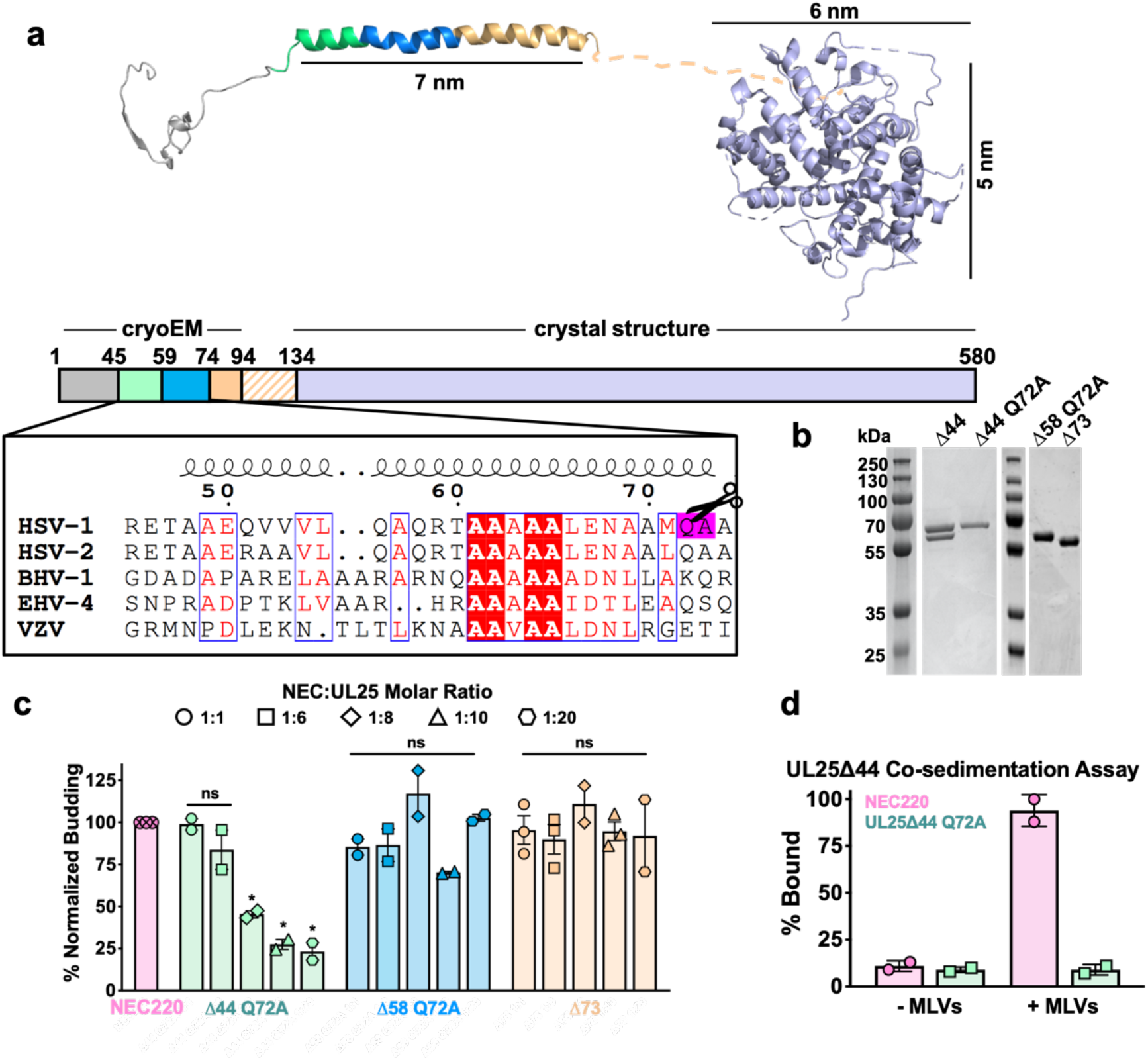
Inhibition of NEC-mediated budding by UL25 constructs. **a)** The UL25 structure and a diagram of domain organization is shown along with a multiple sequence alignment of UL25 residues 45-74 from five alphaherpesviruses. Sequence alignment was generated using Clustal Omega^45^ and displayed using ESPript 3.0^46^. Identical residues are shown as white letters on a red background. Similar residues are shown as red letters in a blue box. Secondary structure derived from the cryoEM reconstruction of HSV-1 UL25 is shown above the alignment. The following herpesvirus sequences were used (GenBank GeneID numbers in parentheses): HSV-1, herpes simplex virus type 1, strain 17 (<u>2703377</u>); HSV-2, herpes simplex virus type 2, strain HG52 (<u>1487309</u>); BHV-1, bovine herpesvirus-1 (<u>4783418</u>); EHV-4, equine herpesvirus-4, strain NS80567 (<u>1487602</u>); and VZV, varicella-zoster virus, strain Dumas (<u>1487687</u>). **b)** SDS-PAGE of purified UL25 constructs: UL25Δ44 (cleaved product; 57 kDa), UL25Δ44 Q72A (single product; 57 kDa), UL25Δ58 Q72A (56 kDa) and UL25Δ73 (54 kDa). **c)** UL25Δ44 Q72A inhibits NEC budding whereas other UL25 constructs do not. For each condition, NEC-mediated budding was tested at 1:1, 1:6, 1:8, 1:10, and 1:20 NEC:UL25 molar ratios. Each construct was tested in at least two biological replicates, consisting of three technical replicates. Symbols show average budding efficiency of each biological replicate relative to NEC220 (100%; pink). Error bars represent the standard error of measurement for at least two individual experiments. Significance compared to NEC220 was calculated using an unpaired t-test against NEC220. *P-value < 0.1. **d)** UL25Δ44 Q72A does not bind to acidic lipid membranes.

### UL25Δ44 Q72A inhibits NEC-mediated budding

To assess the effect of UL25 on NEC-mediated budding, we used an established *in-vitro* budding assay utilizing recombinant, soluble NEC220 (full-length UL31 and UL34 residues 1-220), fluorescently labelled giant unilamellar vesicles (GUVs), and membrane-impermeable fluorescent dye, Cascade Blue (Bigalke et al. 2014). NEC220 and UL25Δ44 Q72A were added to the GUVs in 1:1, 1:6, 1:8, 1:10, or 1:20 molar ratios, and budding events were quantified. UL25Δ44 Q72A inhibited NEC-mediated budding in a dose-dependent manner, and at 1:10 or 1:20 NEC:UL25 molar ratios, few budding events were observed (Fig. 1c). By contrast, UL25Δ73 did not inhibit budding even at a 1:20 ratio of NEC:UL25 (Fig. 1c), which suggested that residues 45-73 were necessary for inhibition.

UL25Δ44 consists of a long N-terminal α-helix (residues 48-94), followed by a flexible linker unresolved in the cryoEM structure, and a C-terminal globular core (residues 134-580) (Fig. 1a). Residues 45-73 encompass the N-terminal half of the long α-helix. To further narrow down the inhibitory region within UL25, we analyzed its sequence conservation. Sequence alignment of UL25 homologs from five alphaherpesviruses revealed a divergent N terminus followed by a highly conserved alanine-rich region, residues 61-69 (Fig. 1a). We generated the UL25Δ58 Q72A construct lacking the divergent N terminus of the α-helix (Fig. 1b). UL25Δ58 Q72A did not inhibit NEC220 budding (Fig. 1c). We also generated a UL25Δ50 Q72A construct (as a control for studies using eGFP-UL25 chimera described below), which inhibited budding at a 1:10 NEC:UL25 ratio, the minimal UL25 concentration for budding inhibition (Fig. 2a). Thus, residues 51-73 appear essential for inhibition whereas residues 45-50 are dispensable.

**Fig. 2.**
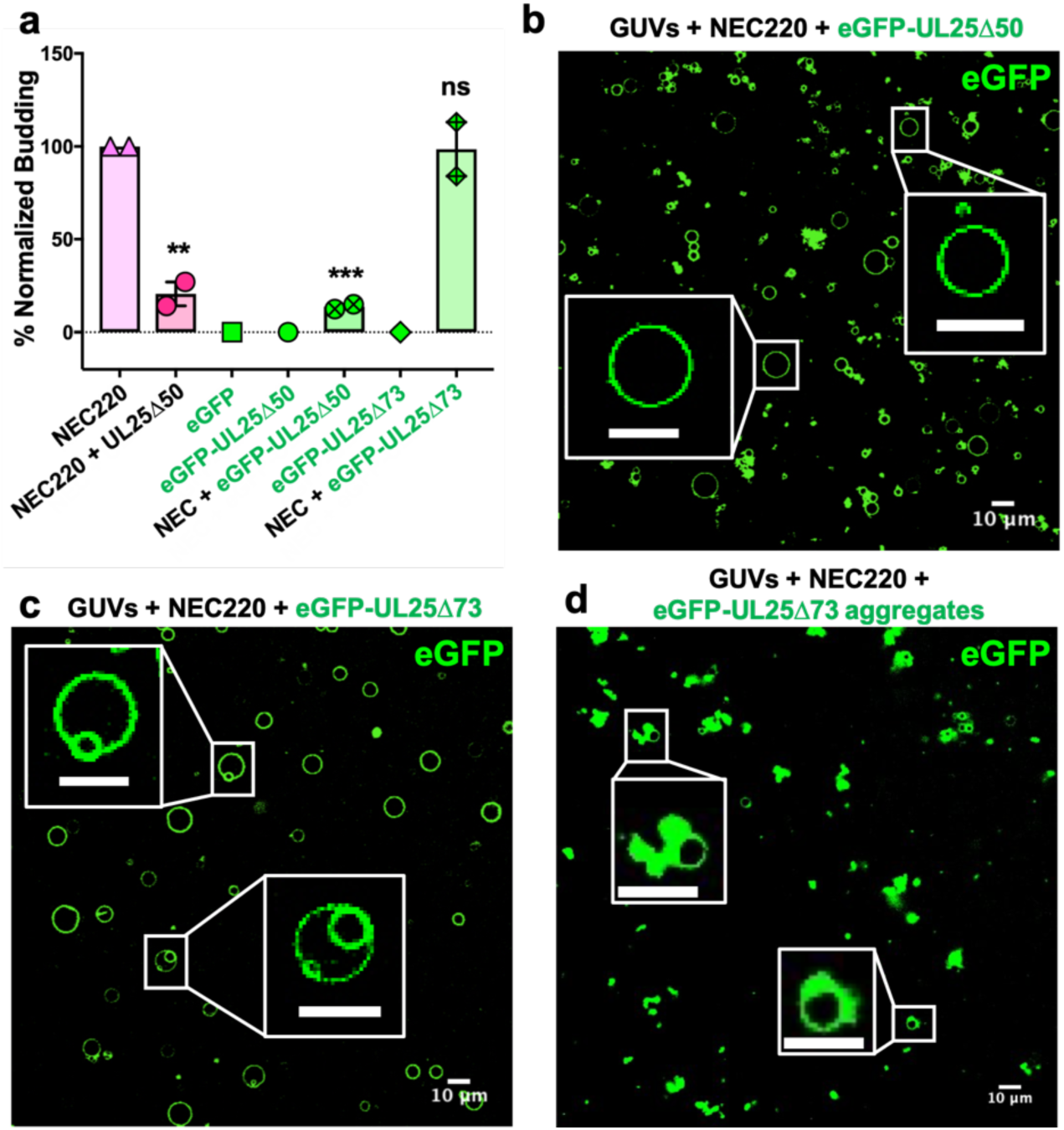
eGFP-UL25Δ50 inhibits NEC budding while eGFP-UL25Δ73 does not. **a)** Quantification of NEC budding in the presence of either eGFP-UL25Δ50 or eGFP-UL25Δ73. Each construct (except in the absence of NEC220) was tested in at least two biological replicates, each consisting of three technical replicates. Symbols show the average budding efficiency of each biological replicate relative to NEC220 (100%). Error bars represent the standard error of measurement for at least two individual experiments. Significance compared to NEC220 was calculated using an unpaired t-test against NEC220. **P-value < 0.01 and ***P-value < 0.001. **b)** Confocal image of eGFP-UL25Δ50 bound to NEC-coated vesicles. No budding is observed. **c)** Confocal image of eGFP-UL25Δ73 either bound to or budded into vesicles with the NEC. **d)** Confocal image of eGFP-UL25Δ73 aggregating on the surface of NEC-coated vesicles. All scale bars = 10 μm.

### UL25 does not bind synthetic membranes

We first tested whether UL25 inhibited NEC-mediated budding by competing with the NEC for binding to membranes. We utilized an established co-sedimentation assay utilizing multilamellar vesicles (MLVs) of the same composition as the GUVs used in the budding assay (Bigalke et al. 2014). Unlike NEC220, UL25Δ44 Q72A did not bind synthetic lipid vesicles (Fig. 1d) and, therefore, could not compete with the NEC220 for binding to membranes.

### UL25Δ44 and NEC do not interact in solution

UL25 does not bind membranes (Fig. 1d) so, to inhibit NEC-mediated budding, UL25 must instead bind to the NEC. However, no binding was detected in solution, either between UL25Δ44 and NEC220 by isothermal titration calorimetry (Supplementary Fig. S1) or between UL25Δ44 and NEC185Δ50 [a truncated construct that was crystallized previously (Bigalke and Heldwein 2015)] by size-exclusion chromatography (Supplementary Fig. S1). Therefore, to bind UL25, NEC may need to be bound to the membrane. Surface plasmon resonance experiments were also performed, but significant nonspecific binding precluded clear data interpretation (data not shown).

### *Both inhibitory UL25*Δ*44 Q72A and non-inhibitory UL25*Δ*73 colocalize with membranes in the presence of the NEC*

To visualize UL25 localization in the presence of NEC and membranes by confocal microscopy, we generated the eGFP-tagged versions of the inhibitory and non-inhibitory UL25 constructs, eGFP-UL25Δ44 Q72A and eGFP-UL25Δ73. However, eGFP-UL25Δ44 Q72A construct was unstable during purification, so eGFP-UL25Δ50 Q72A was generated instead. eGFP-UL25Δ50 Q72A (as well as its untagged version UL25Δ50 Q72A) efficiently inhibited NEC-mediated budding whereas eGFP-UL25Δ73 did not (Fig. 2a). When eGFP-tagged UL25 constructs were incubated with the fluorescently labelled GUVs, no eGFP signal was detected on the GUV membranes (data not shown), confirming that UL25 did not bind membranes directly.

Next, eGFP-UL25Δ50 was incubated with the GUVs in the presence of the NEC220, at a 1:10 molar ratio of NEC to UL25 (the minimal inhibitory UL25 concentration). In the presence of the NEC220, the eGFP-UL25Δ50 colocalized with the GUV membranes, and very little budding was detected (Fig. 2b). UL25 itself does not bind membranes, so instead it must be binding NEC that is bound to the surface of the GUVs.

In the presence of the NEC220, eGFP-UL25Δ73 also colocalized with the GUV membranes. In this case, the eGFP signal was sometimes detected on the membranes of intraluminal vesicles (ILVs) inside the GUVs (Fig. 2c) – a product of budding – which confirmed that binding of eGFP-UL25Δ73 to the NEC220 did not interfere with budding (Fig. 2a) and that eGFP-UL25Δ73 could even remain bound to the NEC-coated membranes throughout budding.

In many cases, however, the eGFP-UL25Δ73 was clustered around the unbudded GUVs, probably due to its aggregation (Fig. 2d). Such aggregation was not observed for eGFP-UL25Δ50 Q72A (Fig. 2b). It is conceivable that the absence of half of the long N-terminal helix of UL25 (Fig. 1a) leads to aggregation of eGFP-UL25Δ73 on NEC-coated GUVs. Although such aggregation inhibits budding locally (Fig. 2d), bulk measurements show that NEC-mediated budding remains efficient in the presence of eGFP-UL25Δ73 (Fig. 2a). We hypothesize that sequestration of large amounts of aggregated eGFP-UL25Δ73 on a few NEC-coated GUVs reduces its concentration throughout the sample, allowing budding to proceed. Taken together, these results suggested that while both inhibitory and non-inhibitory UL25 constructs could bind the membrane-bound NEC, the binding of the inhibitory UL25 construct blocked NEC-mediated budding whereas the binding of the non-inhibitory UL25 construct did not interfere with it.

### Mutations within the putative capsid-binding site on the NEC obviate UL25 inhibition

Residues D275, K279, and D282 at the membrane-distal tip of UL31 have been implicated in capsid binding in PRV (Ronfeldt et al. 2017) and in HSV-1 (Takeshima et al. 2019). We generated a quadruple UL31 mutant in which D275, K279, D282, and a nearby C278 were replaced with alanines. The corresponding mutant NEC220, termed capsid-binding mutant (NEC220-CBM), mediated budding at levels similar to the WT NEC220 (Fig. 3a) but was insensitive to inhibition by UL25Δ44 Q72A (Fig. 3a). Moreover, eGFP-UL25Δ50 Q72A did not co-localize with the GUV membranes in the presence of the NEC220-CBM (Fig. 3b). These results suggested that UL25Δ44 Q72A bound to the membrane-distal tip of UL31 and that this interaction was essential for its inhibitory activity.

**Fig. 3.**
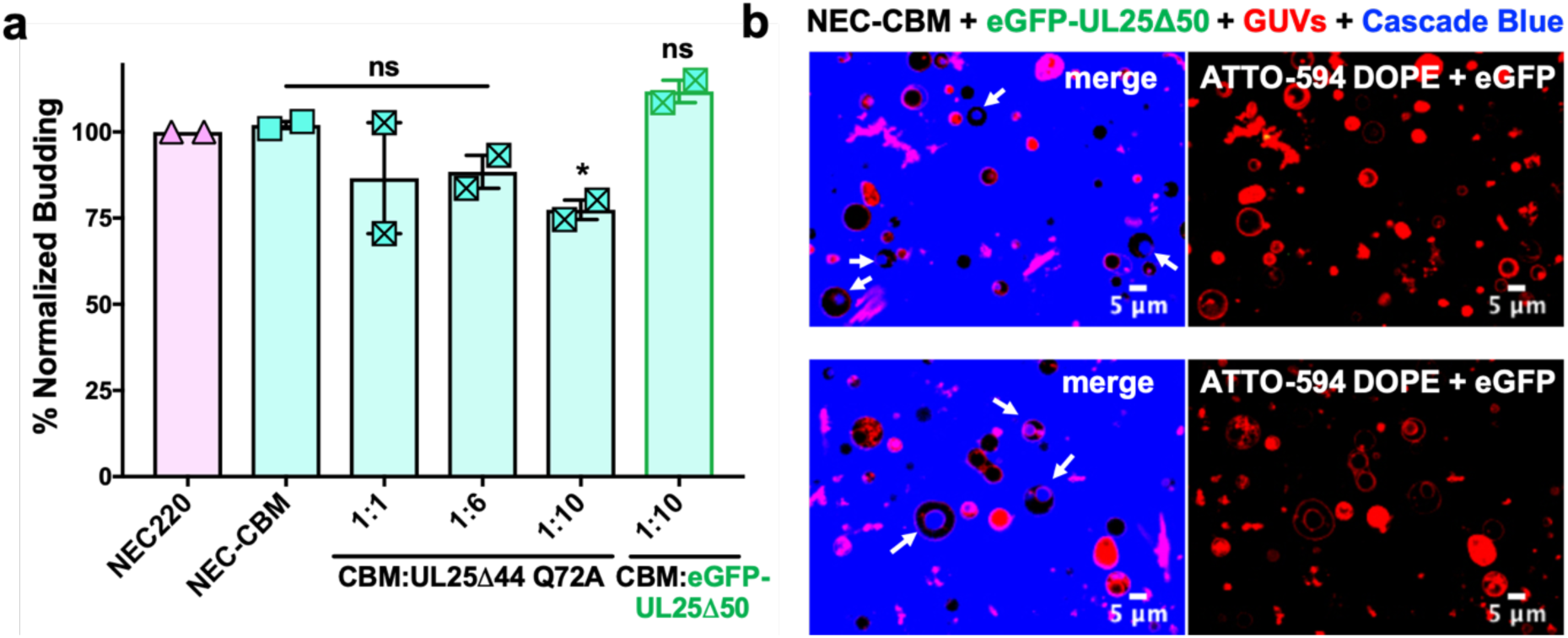
UL25 does not inhibit NEC-CBM budding. **a)** NEC-CBM budding is not inhibited by either UL25Δ44 Q72A or eGFP-UL25Δ50 Q72A. Budding was tested at 1:1, 1:6 and 1:10 NEC220-CBM:UL25 molar ratios for UL25Δ44 Q72A and at a 1:10 NEC-CBM:UL25 molar ratio for eGFP-UL25Δ50 Q72A. Each condition was tested in at least two biological replicates, each consisting of three technical replicates. Symbols represent average budding efficiency of each biological replicate relative to NEC220 (100%). Error bars represent the standard error of measurement for at least two individual experiments. Significance compared to NEC220 was calculated using an unpaired t-test against NEC220. *P-value < 0.1. **b)** Confocal microscopy images showing eGFP-UL25Δ50 Q72A does not bind to NEC-CBM coated GUVs. Intraluminal vesicles due to NEC-CBM budding are indicated by white arrows. Left column is merged with the red (ATTO-594 DOPE), green (eGFP), and blue channels (Cascade Blue). Right column is red (ATTO-594 DOPE) and green (eGFP) only.

### UL25 binds membrane-bound NEC

To understand how UL25 inhibits NEC-mediated budding, we turned to cryoEM. Previously, we showed that NEC-mediated budding of synthetic large unilamellar vesicles (LUVs) resulted in the formation of smaller vesicles containing ∼11-nm thick internal NEC coats (Bigalke et al. 2014). Here, UL25Δ44 Q72A and NEC220 (at a 1:10 molar ratio of NEC to UL25) were incubated with LUVs of the same composition as the GUVs used in the budding assay and visualized by cryoEM. In the presence of UL25Δ44 Q72A and NEC220, the LUVs were mostly spherical, and their external surface was coated with ∼17-nm thick coats (Fig. 4a) although these typically did not cover the entire surface (Fig. 4a and Supplementary Fig. S2). The external coats formed in the presence of UL25Δ44 Q72A are ∼6-nm thicker than the internal NEC coats, and the diameter of the globular portion of UL25 is also ∼ 6 nm. Therefore, we hypothesize that the external coats are composed of a UL25Δ44 Q72A layer positioned on top of the membrane-bound NEC220 layer (Fig. 4a). Very few budded vesicles were observed under these conditions, which is consistent with the inefficient budding observed by confocal microscopy (Fig. 1c). Thus, binding of UL25Δ44 Q72A to the NEC220 on the surface of the lipid vesicles correlated with its ability to inhibit NEC-mediated budding.

**Fig. 4.**
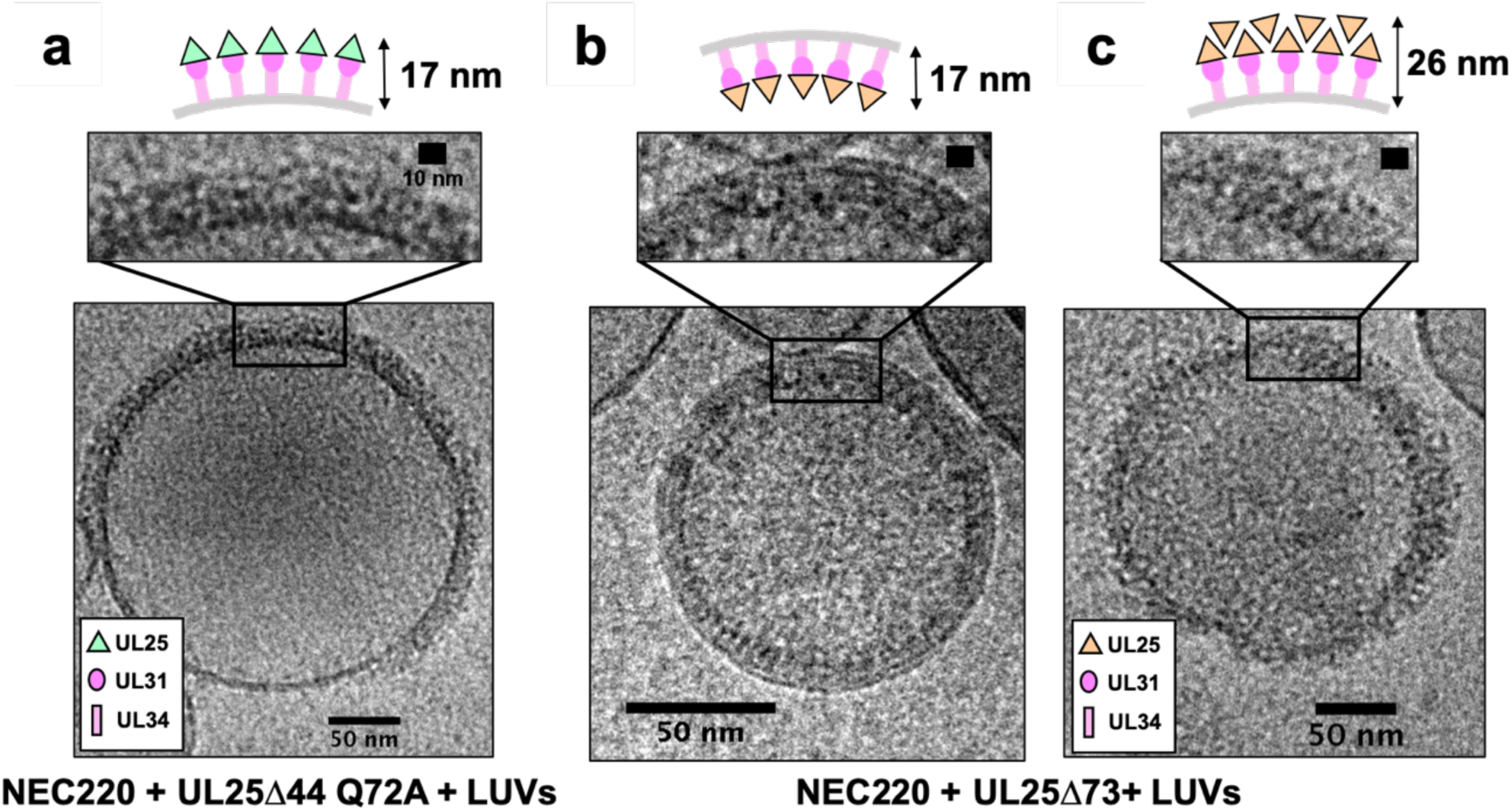
CryoEM shows UL25Δ44 Q72A inhibits NEC budding while UL25Δ73 does not. CryoEM images of NEC-mediated budding in the presence of either UL25Δ44 Q72A **(a)** or UL25Δ73 **(b and c)**. Aggregation of UL25Δ73 is shown in panel C. Scale bars = 50 nm. Inset scale bars = 10 nm.

Co-incubation of UL25Δ73 and NEC220 with LUVs yielded budded vesicles (Fig. 4b) some of which contained ∼17-nm thick internal coats (Fig. 4b), presumably containing UL25Δ73 bound to the NEC220, whereas others contained ∼11-nm thick internal coats (data not shown), presumably containing only NEC220 (Bigalke et al. 2014). We also observed unbudded LUVs containing >25-nm thick heterogeneous protein aggregates on the external surface (Fig. 4c), similar to UL25Δ73 aggregates observed by confocal microscopy (Fig. 2d).

### UL25Δ44 Q72A forms a net of stars bound to NEC pentagons

Interactions between UL25Δ44 Q72A and membrane-bound NEC220 were visualized in three dimensions by cryoET (Fig. 5). Sub-tomographic averaging of the 3D reconstructions of unbudded LUVs coated with NEC220 and UL25Δ44 Q72A (Fig. 5a) revealed that UL25Δ44 Q72A formed a net of five-pointed stars (Fig. 5c) covering the surface of membrane-bound NEC220 whereas the NEC220 formed pentagons (Fig. 5d). Five-pointed stars of UL25 were positioned directly on top of the NEC pentagons (Figs. 5c, d). The star net of UL25 appears to “lock” the NEC layer in place, which could prevent it from undergoing conformational rearrangements required for membrane deformation and budding.

**Fig. 5.**
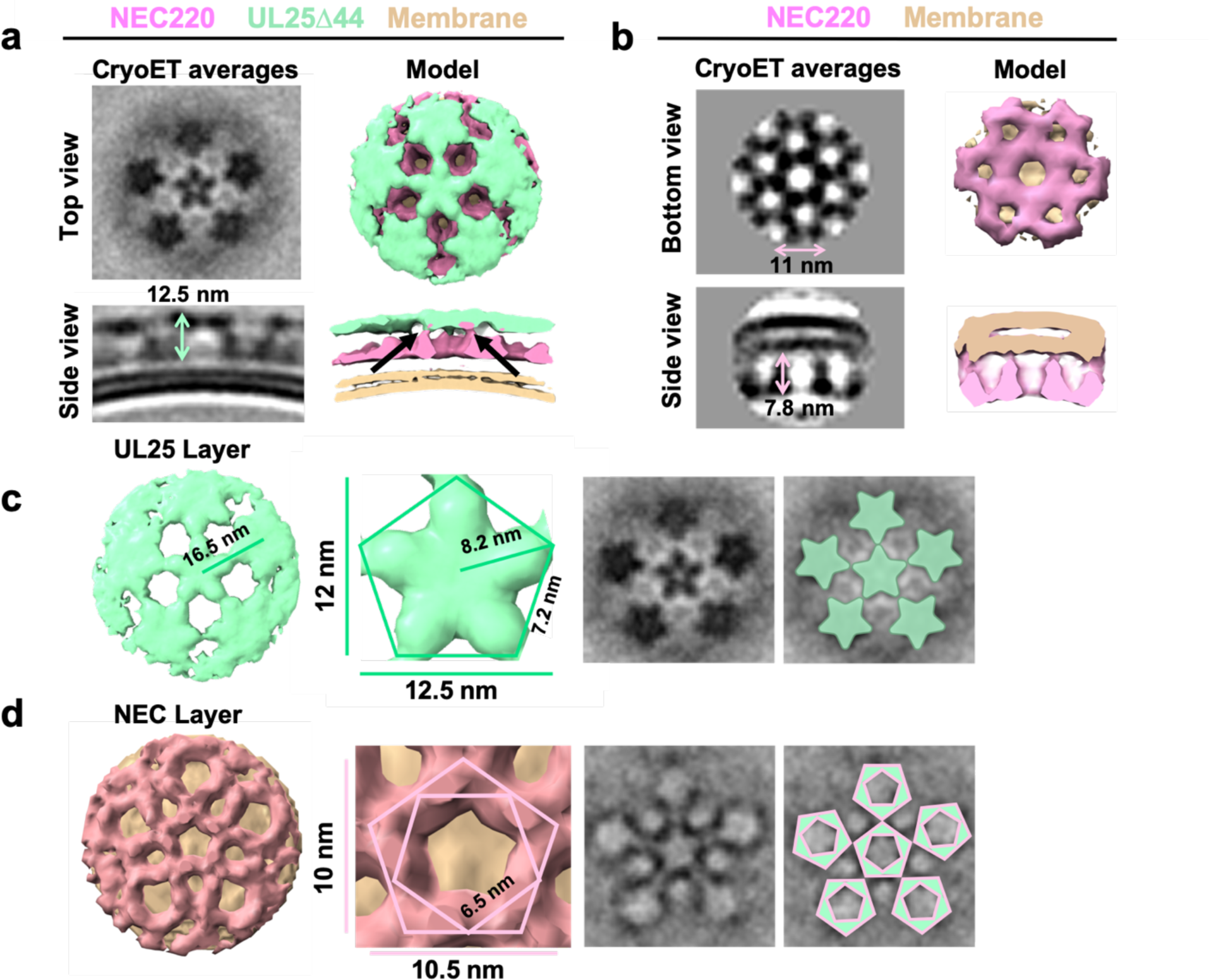
CryoET of UL25-mediated inhibition of NEC budding. **a)** CryoET averages of NEC in the presence of UL25Δ44 Q72A (top and side views). Corresponding 3D models are shown with NEC (pink) and UL25Δ44 Q72A (green). The vesicle bilayer is shown in beige. The models show the UL25 layer coating the NEC layer in five-pointed stars on the outside of the vesicles. The length of the NEC-UL25 spikes is 12.5 nm. Black arrows indicate the point of tilt within the NEC layer. **b)** CryoET averages of NEC forming hexameric lattices in the presence of membranes (bottom and side views). Corresponding 3D models are shown with NEC (pink) and the vesicle bilayer (beige). The diameter of the hexameric rings is ∼11 nm, while the length of the spikes is 7.8 nm. **c)** CryoET model and averages of the UL25 layer (green) highlighting the five-pointed star formation of UL25 (represented here as a pentamer of dimers) in the presence of NEC. **d)** CryoET model and averages of the NEC layer showing NEC forms a pentagonal lattice (pink pentagons), rather than hexagonal (as seen for wild-type in panel b). Green triangles indicate location of UL25 binding to the NEC.

The NEC forms hexagonal coats on budded vesicles formed *in-vitro* (Fig. 5b) (Bigalke et al. 2014) and on perinuclear vesicles formed *in vivo* in NEC-expressing uninfected cells (Hagen et al. 2015), so ability of the NEC to form pentagons was unexpected. The NEC pentagons and hexagons have similar dimensions, ∼10.5 nm vs. ∼11 nm in width (Fig. 5b,d) with ∼6.5 nm vs. ∼6.3 nm sides (Fig. 5d) (Bigalke et al. 2014). We know that the hexagons are hexamers of the NEC heterodimers (Bigalke and Heldwein 2015). Therefore, we hypothesize that the pentagons are pentamers of the NEC heterodimers. The NEC heterodimers within the pentagons appear slightly tilted relative to the plane of the membrane (Fig. 5a) whereas the hexagons are perpendicular to the membrane (Fig. 5b).

It should be noted that an entirely pentagonal lattice would yield a small spherical object with high curvature and icosahedral symmetry – neither of which were observed in our cryoET averages. Given that the NEC/UL25 spikes did not fully coat the vesicles (Fig. 4a), this only permitted averaging of local, rather than global, symmetry, providing a snapshot of NEC/UL25 interactions. Furthermore, a mix of both 400 and 800 nm vesicles were used for these experiments, yielding NEC/UL25-bound vesicles of different sizes, resulting in a difference of curvature upon data averaging, ultimately preventing us from mapping the coordinates of each sub-tomogram back onto the raw data to address this issue. Nevertheless, the cryoET data clearly show the ability of the NEC to form an alternative, pentagonal arrangement in the presence of UL25. Our results document the ability of the NEC to form different oligomers.

## DISCUSSION

The intrinsic ability of the NEC to deform and bud membranes and to oligomerize into a hexagonal coat is well established [reviewed in (Bigalke and Heldwein 2016, Mettenleiter 2016, Bigalke and Heldwein 2017, Roller and Baines 2017)]. However, it is unclear how the capsid triggers the formation of the NEC coat around it or how the NEC coat is anchored to the capsid. Moreover, a purely hexagonal lattice is flat, so it remains unknown how the curvature is generated within the hexagonal NEC coat. Here, we have shown that the NEC forms pentagons when bound to star-shaped UL25 oligomers. We hypothesize that UL25/NEC interactions observed *in vitro* mimic UL25/NEC interactions at the capsid vertices and that NEC pentagons anchor the coat to the capsid. We further hypothesize that NEC pentagons formed at the points of contact with the capsid vertices could nucleate the assembly of and introduce curvature into the hexagonal NEC coats.

### *UL25 inhibits NEC-mediated budding* in vitro *by forming a star-shaped net over the membrane-bound NEC layer*

The inhibitory UL25Δ44 Q72A construct, which is composed of a globular core and a long N-terminal helix, formed five-pointed stars linked into a net on the surface of the membrane-bound NEC220 layer. The five-pointed stars formed by UL25 in our cryoET reconstructions resemble the five-pointed stars that crown each capsid vertex and are composed of five copies of the capsid-associated tegument complex (CATC) (Dai and Zhou 2018) (Fig. 6a). Each CATC is composed of two copies of UL25, one copy of UL17, and two copies of the C-terminal portion of the tegument protein UL36 (Dai and Zhou 2018) and has a characteristic antiparallel four-helix bundle composed of two UL25 helices and two UL36 helices (Fig. 6a).

**Fig. 6.**
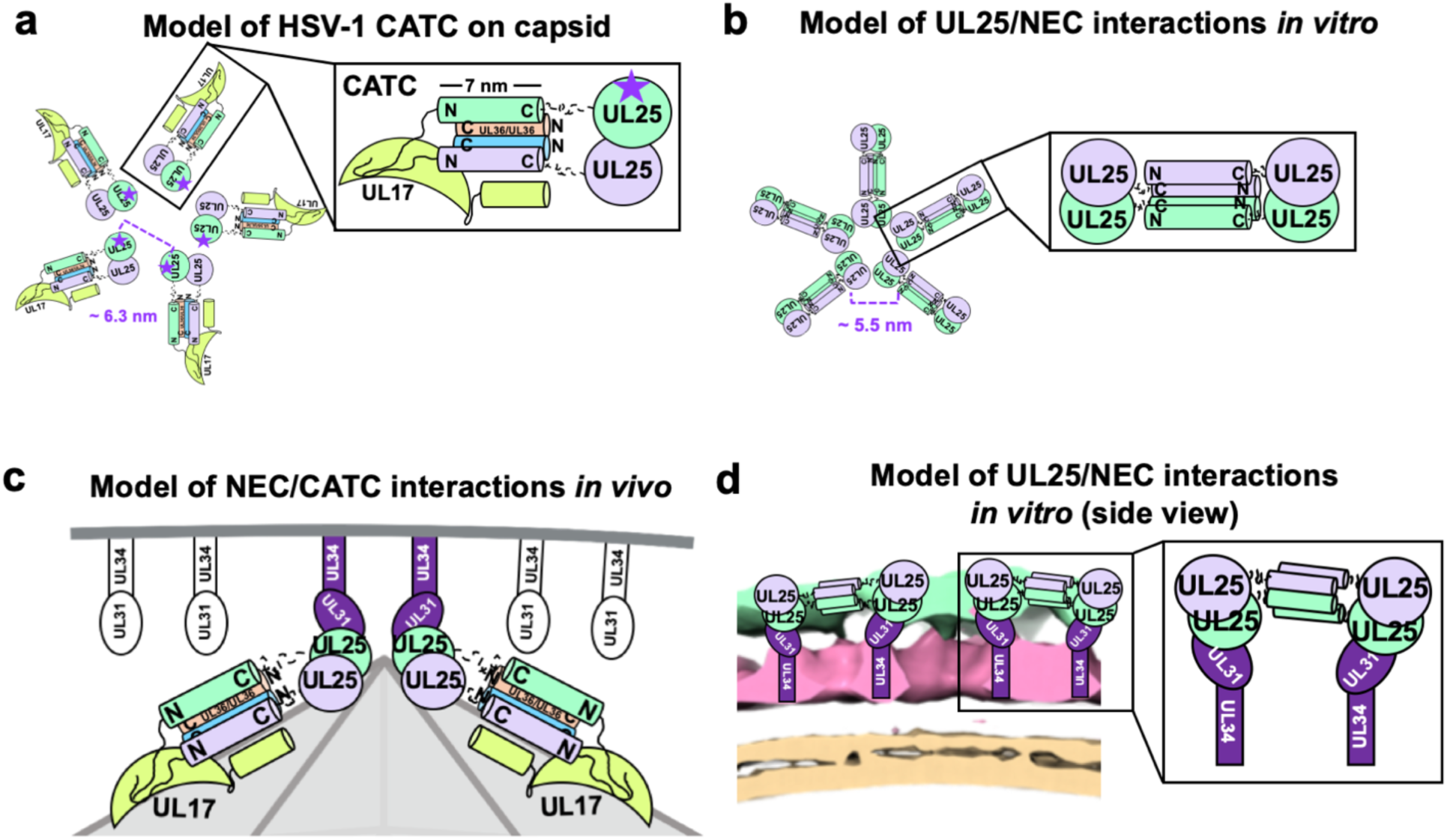
Models of UL25/UL25 and UL25/NEC interactions *in vitro* and *in vivo*. a) A schematic representation of the pentagonal HSV-1 CATC [two copies of UL25 (green and purple), two copies of C-terminal UL36 (peach and blue) and one copy of UL17 (lime green)] arrangement at the capsid vertex. Inset shows a close-up view of the characteristic antiparallel four-helix bundle composed of two UL25 helices and two UL36 helices. Purple stars indicate the proposed UL25 copies that bind to the NEC upon capsid docking. The distance between the centers of two adjacent inner UL25 cores (green) in the capsid (Dai & Zhou, 2018) is ∼6.3 nm. b) Proposed model of the UL25 stars formed *in vitro*. The distance between the centers of two adjacent UL25 dimers is ∼5.5 nm. Inset shows a close-up view of the proposed antiparallel four-helix bundle composed of two pairs of UL25 helices from adjacent stars. We hypothesize that four-helix bundles link the neighboring UL25 stars into a net. c) Proposed side-view model of the NEC (purple) interacting with the most surface exposed capsid-bound UL25 (green), resulting in a pentameric NEC (indicated by dark purple coloring). NEC molecules prior to capsid binding are shown in an unknown oligomeric state (white). d) Side view of the proposed NEC/UL25 interactions *in vitro*.

**Fig. 6.**
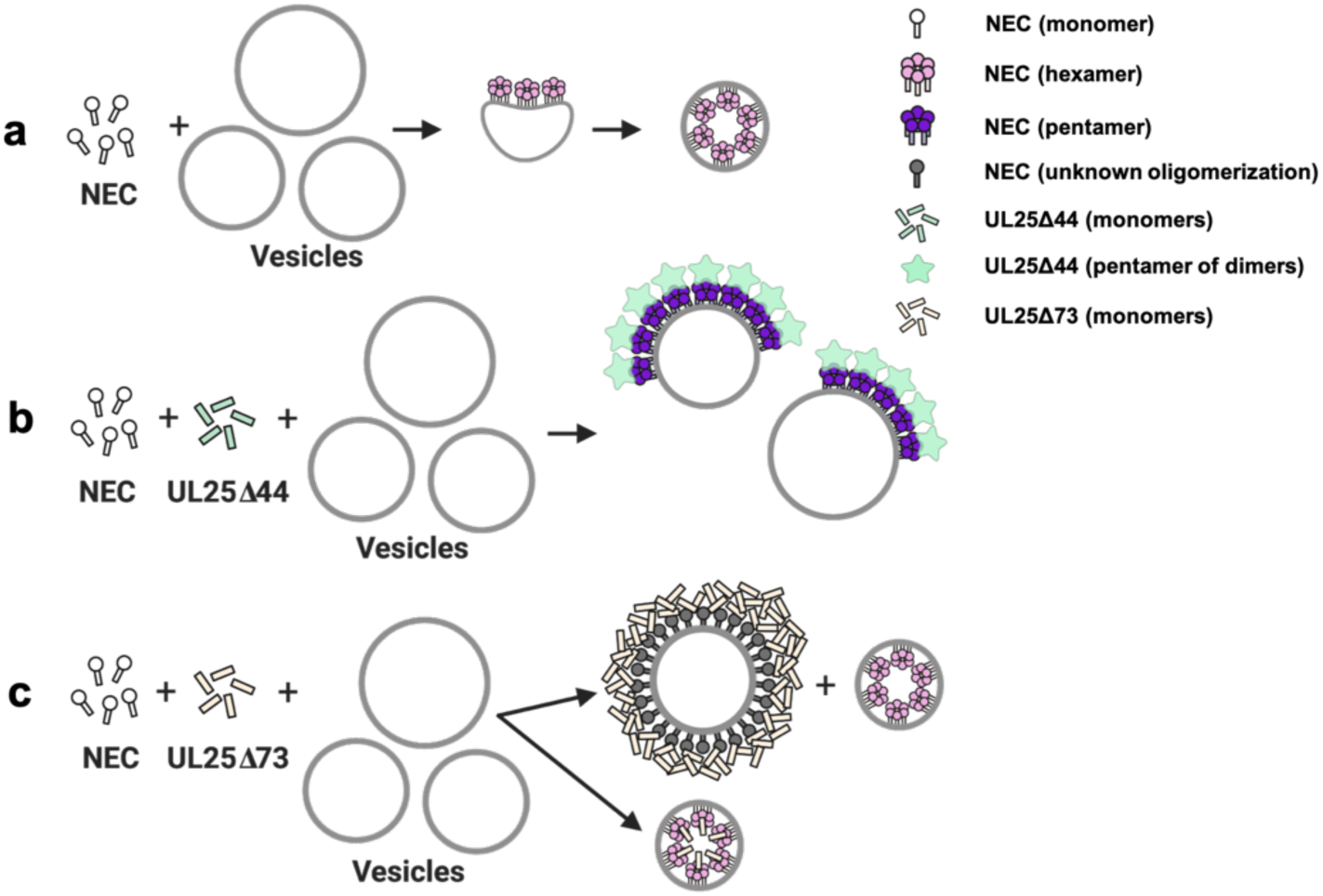
A model of NEC-mediated budding in the absence and presence of UL25, in vitro. **a)** NEC-mediated budding requires only the NEC, which vesiculates membranes by forming hexagonal coats (pink) that, potentially, contain irregular defects to achieve curvature. **b)** UL25Δ44 Q72A (green) inhibits NEC-mediated budding by inducing the formation of a pentagonal NEC coat (purple) suboptimal for budding. **c)** UL25Δ73 (peach) aggregates around some NEC-coated vesicles, which blocks budding. Sequestration of UL25Δ73 at a few locations reduces its concentration elsewhere and enables budding. Binding of UL25Δ73 to NEC in the absence of aggregation does not interfere with budding, and bound UL25Δ73 buds into vesicles with the NEC. This figure was created with Biorender.com.

We hypothesize that when bound to the NEC220 on the membrane surface *in vitro*, UL25Δ44 Q72A also forms an antiparallel four-helix bundle. Only in this case, the bundle is composed of two pairs of UL25 helices from adjacent stars (Fig. 6b). We hypothesize that four-helix bundles link the neighboring UL25 stars into a net. This arrangement of UL25 would require that each UL25 “star” consist of 10 copies of UL25, with cores arranged in the center and 5 pairs of helices radiating out (Fig. 6b). The UL25 cores bind the NEC (Fig. 6d), consistent with previous studies (Yang et al. 2014).

Based on our observations, we propose the following model of NEC-mediated budding *in vitro* and its inhibition by UL25 (Fig. 7). *In vitro*, the NEC-mediated membrane budding leads to the formation of negative membrane curvature and the internal NEC coats on the budded vesicles (Fig. 7a). UL25Δ44 Q72A binds the membrane-bound NEC and forms five-pointed stars on top of NEC pentagons (Fig. 7b) that are linked into a net. Formation of this net could inhibit budding by restricting conformational changes within the NEC lattice necessary to generate negative membrane curvature. By contrast, UL25Δ73 construct does not inhibit budding. Residues 45-73 form about half of the 7-nm-long N-terminal helix (Fig. 1a). Their removal probably precludes formation of stable four-helix bundles. UL25Δ73 is also prone to aggregation likely because the shorter helix is less stable (Fig. 7c).

**Fig. 7.**
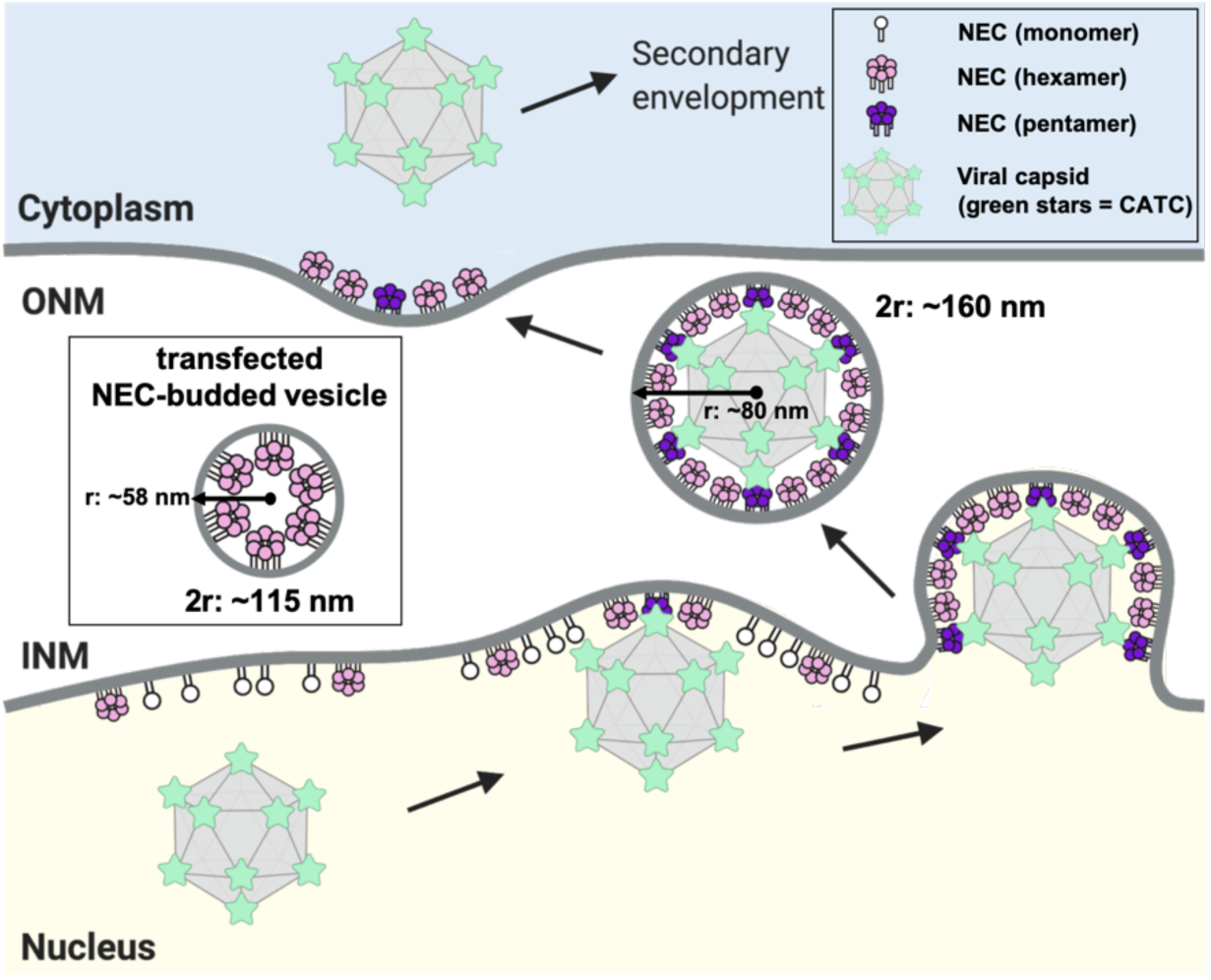
A model of NEC-mediated budding in HSV-1 infected cells. Capsid-bound UL25 induces the formation of pentagonal insertions (purple pentamers) within the NEC coat (pink hexamers and white monomers) as it is forming, which enables the formation of an NEC coat of appropriate size and curvature around the capsid. Inset shows a transfected NEC-budded vesicle which forms a hexagonal coat with presumably irregular defects, similar to the NEC coat formed in vitro. This figure was created with Biorender.com.

### *Inhibition of NEC-mediated budding by UL25 is likely an* in-vitro *phenomenon*

The robust budding ability of the NEC, which is observed both *in vitro* or in NEC-expressing uninfected cells, must be controlled during infection to ensure budding of only mature capsids and to prevent premature, non-productive budding. NEC budding is presumably negatively regulated by a viral protein. It is tempting to speculate that UL25 could inhibit the budding activity of the NEC not only *in vitro* but also during infection. Free UL25 is likely present within the nucleus, but if it were to inhibit NEC-mediated budding, it would then be expected to bind the accumulate at the nuclear rim and accumulate there. Yet, such accumulation has not yet been observed. Therefore, inhibition of NEC-mediated budding by UL25 is likely an *in vitro* phenomenon. Instead, we hypothesize that interactions between UL25 and membrane-bound NEC, which we observed by cryoET and which likely result in budding inhibition *in vitro*, mimic interactions of the NEC with UL25 on the capsid vertices.

### *UL25/NEC interactions* in vitro *mimic interactions between CATC at the capsid vertices and the NEC coats during infection*

We observed that when bound to UL25 *in vitro*, the NEC formed pentagons, which established that the NEC can oligomerize into both hexamers and pentamers and that the oligomeric state of the NEC was influenced by UL25. We hypothesize that NEC pentagons formed at the points of contact with the capsid vertices help anchor the NEC coat to the capsid.

Each capsid vertex is crowned by a five-pointed star composed of five copies of CATC, with 10 globular cores of UL25 arranged in the center as loosely associated dimers (Fig. 6a) (Dai and Zhou 2018). We hypothesize that binding of the cores of five neighboring CATC copies at each capsid vertex (Figs. 6a, b) to the NEC promotes formation of pentagons that attach the coat to the capsid vertices through increased avidity (Fig. 8). The distance between the presumed locations of the cores within the UL25 stars observed *in vitro* is ∼5.5 nm (Fig. 6b). The distance between the cores of the innermost UL25 copies within the CATC stars is somewhat longer, ∼6.3 nm (Fig. 6a), but the cores could move into most favorable orientations for binding the NEC. The dynamic nature of the UL25 cores is evident from the cryoEM reconstructions, which show the cores connected to the N-terminal helices by long, flexible linkers (Dai and Zhou 2018). On the other hand, the NEC may be able to tilt relative to the membrane surface (Fig. 5a), which would also allow each NEC to adopt the optimal orientation for binding the UL25 cores at the capsid vertices.

UL25 and the NEC – a monomer and a heterodimer, respectively – do not interact in solution but do interact when the NEC is bound to the membrane, which implies the importance of avidity in binding. The oligomeric state of the NEC prior to the arrival of the capsid is unknown, but high concentration of the NEC at the inner nuclear membrane (Hagen et al. 2015, Newcomb et al. 2017) would provide enough NEC copies locally to form a binding site for UL25 at the capsid vertex.

UL25 likely binds to the membrane-distal tip of UL31 because mutations within that region render NEC-CBM mutant insensitive to inhibition by UL25. These results are in agreement with the study by Takeshima *et al.* that showed this UL25-NEC interaction involved UL31 residues R281 and D282 (Takeshima et al. 2019). Other charged residues within the membrane-distal region of UL31 have also been implicated in capsid interactions in PRV (Ronfeldt et al. 2017). Thus, we hypothesize that binding of the capsid to the NEC during nuclear egress are mediated by UL25/UL31 interactions. Indeed, NEC185Δ50 (a previously crystallized truncated construct with an intact membrane-distal UL31 region) from *E. coli* binds purified nucleocapsids from HSV-1 infected cells, and this interaction requires UL25 rather than the VP5, VP23, or UL17 capsid proteins (Takeshima et al. 2019). Nucleocapsids can also bind free UL31 (Yang and Baines 2011, Yang et al. 2014).

### How does the hexagonal NEC lattice achieve curvature?

The ability of the NEC to oligomerize into a hexagonal lattice *in vitro* and *in vivo* is well documented (Bigalke et al. 2014, Bigalke and Heldwein 2015, Hagen et al. 2015) and is an important feature of its membrane deformation mechanism (Roller et al. 2010, Bigalke et al. 2014, Bigalke and Heldwein 2015). But how the hexagonal NEC lattice accommodates curvature is yet unclear.

A strictly hexagonal lattice is flat, so the curvature is typically achieved through the inclusion of lattice defects, also termed insertions. These can either be regular insertions of a different geometry, for example, pentagons – as observed in icosahedral or fullerene-like capsids – or irregular insertions. Although no deviations from the hexagonal symmetry have yet been visualized in any NEC coats (Bigalke et al. 2014, Hagen et al. 2015, Newcomb et al. 2017), this could be due to the low resolution of the cryoET reconstructions or the imposition of symmetry in averaging. For example, one study used cryoEM and cryoET to visualize NEC/capsid interactions within perinuclear enveloped virions isolated from cells infected with an US3 kinase-null HSV-1 (Newcomb et al. 2017), a mutation that causes the accumulation of perinuclear enveloped virions. Although only the hexagonal NEC arrays were observed, the averaging of NEC/CATC interactions was hindered by significant noise in the relevant regions of the tomograms because the NEC coat did not have the same icosahedral symmetry as the capsid. As the result, NEC/CATC interactions were difficult to visualize. Additionally, the lack of US3 could have altered the structure of the NEC coat or the NEC/CATC interactions.

In perinuclear viral particles formed in PRV-infected cells, the NEC coats appear tightly associated with the capsid (Hagen et al. 2015). Capsidless perinuclear vesicles formed in uninfected cells expressing PRV NEC (Hagen et al. 2015) are relatively uniform in size (∼115 nm in diameter; Fig. 8, inset) but smaller than the capsid (∼125 nm in diameter (Liu et al. 2017, Dai and Zhou 2018)) or the perinuclear vesicles isolated from cells infected with the HSV-1 US3-null mutant virus (∼160 nm in diameter (Newcomb et al. 2017)) (Fig. 8). The capsid thus appears to define the size of NEC-budded vesicles during infection, so the capsid geometry could influence the geometry of the NEC coat.

Based on our observation that the NEC forms pentagons when bound to UL25, we hypothesize that during nuclear egress, NEC pentagons formed at the points of contact with the capsid vertices not only anchor the NEC coat to the capsid but also generate NEC coat of appropriate curvature through the inclusion of pentagons into a hexagonal coat as it assembles around the capsid. A similar strategy is observed during HIV-1 capsid formation by the Gag protein (Briggs et al. 2009, Schur et al. 2015). As the mature capsid is built, the Gag protein is cleaved, and the Gag capsid-domain builds a hexagonal lattice containing 12 pentamers to form a closed fullerene-like structure.

We do not yet understand how curved NEC coats are assembled in the absence of capsid. Hexagonal NEC coats formed in *in vitro* or in NEC-expressing cells have a smaller diameter than those formed around the capsid (Fig. 8), so they may achieve coat curvature by other means, for example, by having irregular defects. Incorporation of irregular defects into curved hexagonal lattices have been observed for immature HIV capsids formed by Gag protein (Briggs et al. 2009, Schur et al. 2015) and in early poxvirus envelopes formed by the D13 protein (Heuser 2005, Hyun et al. 2011). NEC could, potentially, use a similar strategy in the absence of a capsids.

### A model of NEC-mediated capsid budding during infection

Based on our observations, we propose the following model of NEC-mediated capsid budding during nuclear egress in infected cells (Fig. 8). Binding of the cores of five neighboring copies of CATC at the capsid vertices to the NEC at the INM would promote formation of NEC pentagons, which would help anchor the capsid to the INM and could also serve as a nucleation event for the assembly of the NEC coat around the capsid. As the hexagonal NEC coat continues to grow, the incorporation of pentagons into the coat at the points of contact with the vertices would both help attach the NEC coat to the capsid and introduce curvature into the NEC coat (Fig. 8).

In both HSV-1 and PRV, removal of UL25 results in an accumulation of capsids at the INM, unable to undergo egress (Klupp et al. 2006, Kuhn et al. 2008). Our results suggest that UL25 both anchors the NEC coat to the capsid and contributes to formation of a curved coat. Additionally, our results could potentially explain why mostly mature, DNA-containing C-capsids undergo budding at the INM (Roizman and Furlong 1974, Klupp et al. 2011). A- and B-capsids have fewer UL25 copies on the capsid surface (Newcomb et al. 2006), and we hypothesize that only C-capsids, which contain UL25 at a full occupancy, can generate pentagonal NEC insertions necessary for the formation of an NEC coat around the capsid. In this manner, NEC/UL25 interactions could provide a quality-control mechanism that would favor budding of mature, DNA-containing C-capsids – which have a full UL25 set – over the immature capsid forms with fewer UL25 copies thereby acting as a checkpoint during nuclear egress.

## ACKNOWLEDGMENTS

We thank Janna Bigalke for generating the UL25Δ44 and UL25Δ73 plasmids, purifying the corresponding proteins, and performing the ITC and the size-exclusion experiments. We thank Alenka Lovy (Tufts University) for assistance with fluorescence microscopy experiments and Mike Rigney (Brandeis University) for assistance with cryoEM imaging. We also thank Peter Cherepanov (Francis Crick Institute) for the gift of the GST-PreScission protease expression plasmid and Thomas Schwartz (Massachusetts Institute of Technology) for the gift of LoBSTr cells. ITC experiments were performed at the Center for Macromolecular Interactions in the Department of Biological Chemistry and Molecular Pharmacology at Harvard Medical School. CryoEM images were collected at the Electron Microscopy Facility at Brandeis University. CryoET data were collected at the Electron Imaging Center for Nanomachines at the University of California, Los Angeles. This work was funded by the NIH grants R01GM111795 (E.E.H.), 1S10OD018111 (Z.H.Z.), 1U24GM116792 (Z.H.Z.), a Faculty Scholar grant from Howard Hughes Medical Institute (E.E.H.), NIH postdoctoral fellowship F32GM126760 (E.B.D.), the NSF grants DBI-1338135 (Z.H.Z.) and DMR-1548924 (Z.H.Z.), Burroughs Wellcome Fund Collaborative Research Travel Grant (E.B.D.), and the Natalie V. Zucker Research Grant (E.B.D.).

## AUTHOR CONTRIBUTIONS

E.B.D. and E.E.H. designed and coordinated the project; E.B.D. performed the experiments (with the exception of the ITC and size-exclusion experiments) under the guidance of E.E.H; E.B.D. and J.Z. collected cryoET data under the guidance of Z.H.Z; J.Z. processed the cryoET data; all authors analyzed the data, interpreted the results and wrote the manuscript.

## DATA AVAILABILITY STATEMENT

The EM datasets generated in this study will be deposited into the Electron Microscopy Data Bank and will be immediately available upon publication.

## METHODS

### Cloning

All primers used in cloning are listed in Supplementary Table S1. Codon-optimized UL25 gene from HSV-1 strain KOS was synthesized by GeneArt. Digested PCR fragments encoding UL25Δ44 were subcloned by restriction digest into the pJP4 plasmid, which contains a His6-SUMO-PreScission tag in frame with the BamHI restriction site of the multiple-cloning site in a pET24b vector, creating the pJB104 plasmid. DNA fragments encoding UL25Δ50 and UL25Δ73 were amplified by PCR from pJB104 (UL25Δ44) and subcloned into pJP4 by restriction digest using BamHI and XhoI, creating the UL25Δ50 (pED13) and UL25Δ73 (pJB123) plasmids. Site-directed mutagenesis of pJB104 yielded the UL25Δ44 Q72A mutant plasmid (pED03).

DNA encoding the eGFP sequence was PCR amplified out of the eGFP-N2 plasmid (Clontech) and subcloned via single-cut restriction digest into the corresponding UL25 plasmid harboring the cleavable His6-SUMO tag [(either UL25Δ50 (pED13) or UL25Δ73 (pJB123)] creating either the eGFP-UL25Δ50 (pED14) or eGFP-UL25Δ73 (pED05) constructs.

Site-directed mutagenesis of pKH90 (UL31 1-306) using a splicing by overlap extension protocol(Heckman and Pease 2007) followed by restriction digest into the pJP4 vector was used to create the UL31 D275A/C278A/K279A/D282A mutant (pJB118) in the capsid binding mutant construct (NEC-CBM).

### Expression and purification of NEC constructs

Plasmids encoding HSV-1 UL31 1-306 (pKH90) and UL34 1-220 (pJB02) were co-transformed into *Escherichia coli* BL21(DE3) LoBSTr cells (Kerafast) to generate NEC220 (Bigalke et al. 2014). Plasmids encoding HSV-1 UL31 1-306 D275A/C278A/K279A/D282A (pJB118) and UL34 1-220 (pJB02) were co-transformed into *E. coli* BL21(DE3) LoBSTr cells (Kerafast) to generate NEC-CBM. All constructs were expressed using autoinduction at 37 °C in Terrific Broth (TB) supplemented with 100 µg/mL kanamycin, 100 µg/mL ampicillin, 0.2% lactose and 2 mM MgSO4 for 4 h. The temperature was then reduced to 25 °C for 16 h. Cells were harvested at 5,000 x g for 30 min. NEC proteins were purified as previously described(Bigalke et al. 2014) with slight modifications. The NEC220 and NEC-CBM constructs were passed over 2 x 1 mL HiTrap Talon columns (GE Healthcare), rather than ion exchange as previously described, to remove excess cleaved His6-SUMO before injection onto size-exclusion chromatography (as previously described).

### Expression and purification of UL25 constructs

Plasmids encoding either HSV-1 UL25 or eGFP-UL25 constructs were transformed into *E. coli* BL21(DE3) LoBSTr cells and expressed using autoinduction at 37 °C in TB supplemented with 100 µg/mL kanamycin, 0.2% lactose, and 2 mM MgSO4 for 4 h. The temperature was then reduced to 25 °C for 16 h. Cells were harvested at 5,000 x g for 30 min. All purification steps were performed at 4 °C. UL25 constructs were purified in lysis buffer (50 mM Na HEPES pH 7.5, 500 mM NaCl, 1 mM TCEP, and 10% glycerol). Cells were resuspended in lysis buffer supplemented with Complete protease inhibitor (Roche) and lysed with a microfluidizer (Microfluidics). The cell lysate was clarified by centrifugation at 13,000 x g for 35 min and was passed over Ni-NTA sepharose (GE Healthcare) column. The column was subsequently washed with 20 mM and 40 mM imidazole lysis buffer and bound proteins were eluted with 250 mM imidazole lysis buffer. The His6-SUMO tag was cleaved for 16 h using PreScission Protease produced in-house from a GST-PreScission fusion protein expression plasmid. As a final purification step, UL25 constructs were purified with size-exclusion chromatography using either a Superdex 75 or 200 column (GE Healthcare) equilibrated with gel filtration buffer (20 mM Na HEPES, pH 7.0, 100 mM NaCl, and 1 mM TCEP). The UL25 constructs were purified to homogeneity as assessed by 12% SDS-PAGE and Coomassie staining. Fractions containing UL25 were concentrated up to ∼30 mg/mL and stored at −80 °C to prevent degradation observed at 4 °C. Protein concentration was determined by absorbance measurements at 280 nm. The typical yield was 35 mg/L of TB culture.

### Co-sedimentation assay

Co-sedimentation of UL25Δ44 to acidic multilamellar vesicles (MLVs) was performed as previously described (Bigalke et al. 2014). MLVs were prepared in a 3:1:1 ratio of 1-palmitoyl-2-oleoyl-glycero-3-phosphocholine (POPC):1-palmitoyl-2-oleoyl-sn-glycero-3-phospho-L-serine (POPS):1-palmitoyl-2-oleoyl-sn-glycero-3-phosphate (POPA) (Avanti Polar Lipids). Background signal in the absence of liposomes is due to protein aggregation during centrifugation.

### In vitro GUV budding assays

Giant unilamellar vesicles (GUVs; used for their large size and ease of identification at the microscope) were prepared as previously described (Bigalke et al. 2014). For NEC220 only budding quantification, a total of 10 μL of GUVs with a 3:1:1 ratio of POPC:POPS:POPA containing ATTO-594 DOPE (ATTO-TEC GmbH) at a concentration of 0.2 μg/μL was mixed with 1 μM NEC220 (final concentration), and 0.2 mg/mL (final concentration) Cascade Blue Hydrazide (ThermoFisher Scientific). For the NEC and UL25 titration experiments, 10 μL of GUVs and either 1, 6, 8, 10 or 20 μM of UL25Δ44 Q72A, UL25Δ58 Q72A or UL25Δ73 (final concentration) were incubated with 1 μM of NEC220 (final concentration) along with Cascade Blue. For NEC-CBM and UL25 titration experiments, 10 μL of GUVs and either 1, 6, or 10 μM of UL25Δ44 Q72A (final concentration) were incubated with 1 μM of NEC-CBM (final concentration) along with Cascade Blue. The total volume of each sample during imaging for all experiments was brought to 100 μL with gel filtration buffer and the reaction was incubated for 5 min at 20 °C. Samples were imaged in a 96-well chambered cover-glass. Images were acquired using a Nikon A1R Confocal Microscope with a 60x oil immersion lens at the Tufts Imaging Facility in the Center for Neuroscience Research at Tufts University School of Medicine. Images of NEC budding in the presence of eGFP-UL25 constructs were recorded after incubation of 10 μL of GUVs with 10 μM (final concentration) of either eGFP-UL25Δ50 Q72A or eGFP-UL25Δ73 and 1 μM of NEC220 (final concentration). Quantification was performed by counting vesicles in 15 different frames of the sample (∼300 vesicles total). Raw data values for all experiments are listed in Supplementary Table S2. Each condition was tested in at least two biological replicates. Prior to analysis, the background was subtracted from the raw values. The reported values represent the average budding activity relative to NEC220 (100%). The standard error of the mean is reported for each measurement. Significance compared to NEC220 was calculated using an unpaired one-tailed *t*-test against NEC220.

### Isothermal titration calorimetry (ITC)

ITC measurements were recorded using a Microcal ITC200 (Malvern Panalytical) at the Center for Macromolecular Interactions at Harvard Medical School. A solution of UL25Δ44 (200 μM) was titrated into a solution of NEC220 (20 μM) in 20 mM Na HEPES, pH 7.0, 150 mM NaCl, 1 mM TCEP. Control experiments were performed by injecting UL25Δ44 into buffer. Thermograms were plotted by subtracting heats of the control experiments from the sample experiments. The data were not fit due to no detectable binding.

### Cryoelectron microscopy and tomography

A volume of 10 μL of a 1:1 mixture of 400-nm and 800-nm large unilamellar vesicles (LUVs) made of 3:1:1 POPC:POPS:POPA [prepared as previously described (Bigalke et al. 2014)] were mixed on ice with a 30 μL solution of NEC220 and either UL25Δ44 Q72A or UL25Δ73, yielding an NEC:UL25 ratio of 1:10 (NEC concentration was at 1 mg/mL). After 30 min, 3 μL of sample was applied to glow-discharged (30 s) Quantifoil copper grids (R2/2, 200 mesh, Electron Microscopy Sciences), blotted on both sides for 4 s, and vitrified by rapid freezing in liquid ethane (Vitrobot). Grids were stored in liquid nitrogen until loaded into a Tecnai F20 transmission electron microscope (FEI) via a cryo holder (Gatan). The microscope was operated in low dose mode at 200 keV using SerialEM (Mastronarde 2005) and images were recorded with a 4k x 4k charge coupled device camera (Ultrascan, Gatan) at 29,000-fold magnification (pixel size: 0.632 nm). 2D cryo-EM images were recorded at defocus values of −4 to −8 μm and an electron dose ∼15 e/Å^2^. Images are displayed using ImageJ (Schindelin et al. 2015).

For single-axis cryoET data used to generate 3D EM data, samples were incubated on ice for 30 min, and 0.8 μL of 10 nm colloidal gold coated with protein A (Cell Microscopy Core, University Medical Center Utrecht, Department of Cell Biology) was added to the solution and mixed. The mixture (2.5 μL) was applied to freshly glow-discharged (30 s) Quantifoil R 3.5/1 grids (Electron Microscopy Sciences) and manually blotted before being flash-frozen in liquid ethane. Grids were loaded into a FEI Titan Krios electron microscope equipped with a Gatan imaging filter (GIF) and a Gatan K2 summit direct electron detection camera (Roper Technologies, Inc.), operated at 300 kV. The acquisition for automated cryoET tilt series collection was performed using SerialEM (Mastronarde 2005). A tilt series was collected in which the sample was tilted from 0° to +60° degrees and then from 0° to −60°, each in a stepwise fashion with 2° increments. Tilt series were acquired at a magnification of x53,000 (corresponding to a calibrated pixel size of 2.6 Å) with a maintained defocus value of −3 to −4 μm. The total electron dose was ∼100 e/Å^2^.

### 3D reconstruction and subtomographic averaging

The detailed steps of the 3D reconstruction and subtomographic averaging were previously described (Si et al. 2018). Briefly, frames from each recorded tilt series were drift-corrected and averaged with *Motioncorr* (Mastronarde 2005) and was further reconstructed with contrast transfer function (CTF) correction using the IMOD software package (Kremer et al. 1996). Two resulting tomograms were produced by the weighted back projection and simultaneous iterative reconstruction technique (SIRT) methods. A total of 1200 particles were picked for tomograms containing LUVs, NEC220 and UL25Δ44 Q72A. 3D sub-tomographic averaging was completed as described(Si et al. 2018) using the PEET (particle estimation for electron tomography) software (Nicastro et al. 2006). Five-fold symmetry was only applied after five-fold symmetry was apparent in the averaged structure. The original dataset was split into two separate groups, even group and odd group, and averaged independently. Gold standard Fourier Shell Correlation (FSC) analysis for the averaged structure was performed by *calcUnbiasedFSC* in PEET when the two averaged structures converged. The reported resolution is 29 Å based on the 0.143 gold-standard FSC criterion. EM maps will be deposited into the Electron Microscopy Data Bank (EMDB) for immediate access upon publication.

