## Supplemental figures S1-S2, Supplemental Tables S1-S2 for "Structural basis for capsid recruitment and coat formation during HSV-1 nuclear egress"

a

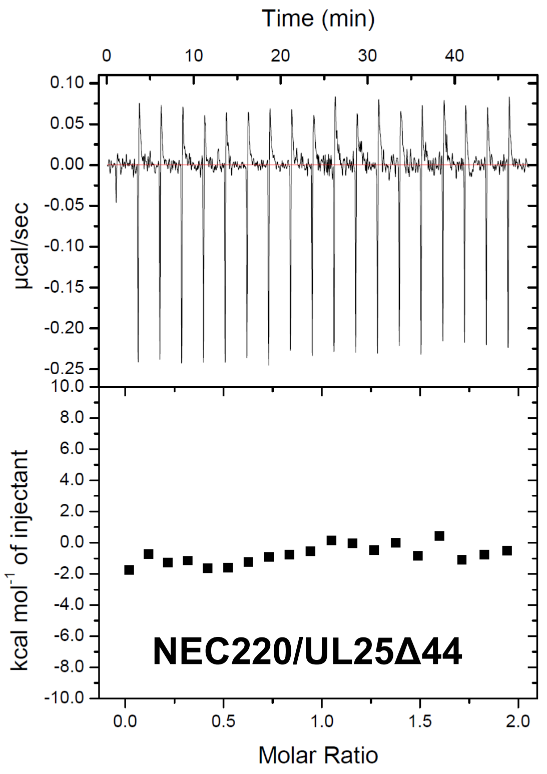


b

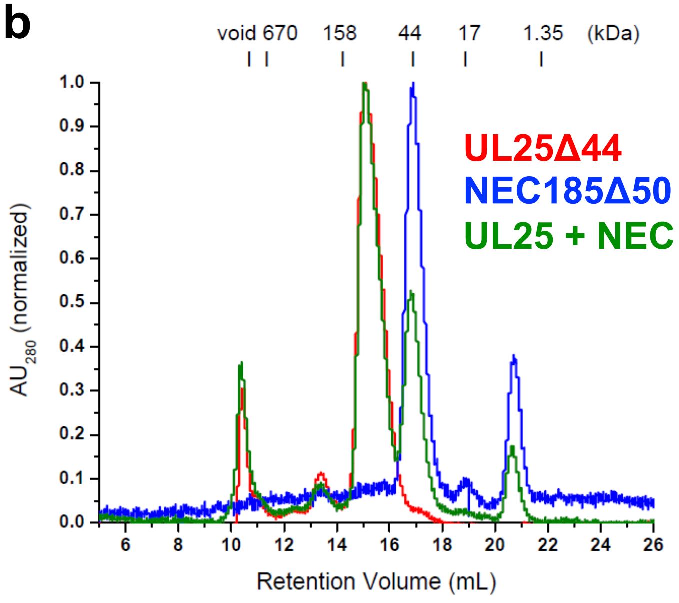


**Supplementary Fig. S1. NEC-UL25 binding studies. a)** ITC of NEC220 and UL25Δ44 showing these two proteins do not bind in solution. **b)** Size-exclusion chromatography of NEC185Δ50 (crystallization construct) and UL25Δ44 shows these proteins also do not bind in solution.

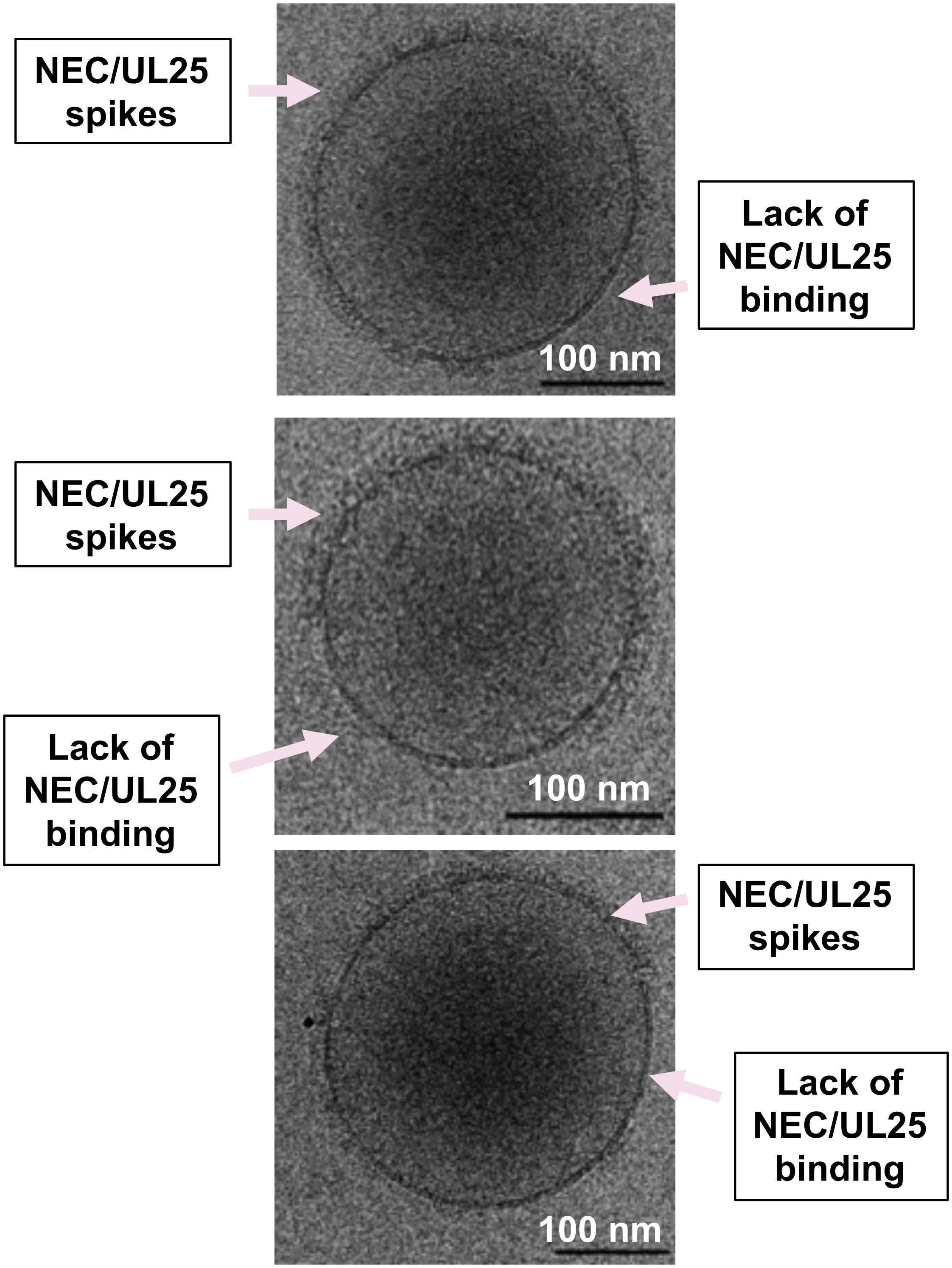


**Supplementary Fig. S2. Incomplete distribution of NEC-UL25 around vesicles.** Slices of selected tomograms of NEC/UL25-bound vesicles used to generate cryoET averages in Figure 5. Regions of either protein binding or lack of protein binding are indicated by pink arrows. Scale bar represents 100 nm.

**Supplementary Table S1. List of primers used for cloning procedures described in Materials and Methods.** All primers are listed in the 5’-3’ direction. Restriction sites are underlined and mutations are bolded.

| **Primer Name** | **Primer Sequence (5’-3’)** | **Restriction Site** |
|---|---|---|
| UL25Δ44 fwd (JB133) | aaaaaaggatcccgtgaaaccgcagcagaacagg | BamHI |
| UL25Δ50 fwd (ED034A) | aaaaaaggatcccaggttgttgttctgcaggcacagcgt | BamHI |
| UL25Δ73 fwd (JB238) | aaaaaaggatccgccgaactgccggttgatattg | BamHI |
| UL25 rev (JB134) | aaaaaactcgagttattacactgcgctcagatactgagg | XhoI |
| UL25Δ44 Q72A (ED006) | aatgcagcaatg**gcg**gcagcc | Site-directed |
| UL25Δ44 Q72A (ED007) | ggctgc**cgc**cattgctgcatt | Site-directed |
| eGFP fwd (ED010) | aaaaaaggatccgtgagcaagggcgaggagctg | BamHI |
| eGFP rev (ED049) | aaaaaaggatcccttgtacagctcgtccatgccgagagtg | BamHI |
| UL31 CBM fwd (JB208) | ccgtgtcggccgca**gcc**atttat**gctgca**atgagg**gcc**atcagcttcgacggg | SOE |
| UL31 CBM rev (JB209) | cccgtcgaagctgat**ggc**cctcat**tgcagc**ataaat**ggc**tgcggccgacacgg | SOE |

**Supplementary Table 2. Raw data and background values collected for each reported GUV budding assay.** Each technical replicate is highlighted by a given color. Technical replicates for each construct collected during the same experiment are highlighted in identical colors within each designated section. The reported biological replicate average values (%) are the points presented in the corresponding manuscript figures.

| | Ratio | ILVs/GUVs counted | Background (BG) | Raw Value (ILV-BG) | Average (AVG) | WT (ILVs/GUVs counted) | WT - BG | WT AVG | % Normalized Budding (Raw Value/WT AVG) * 100 | Biological Replicate Average (%) |
|---|---|---|---|---|---|---|---|---|---|---|
| **UL25 Truncation Constructs** | | | | | | | | | | |
| **NEC220 + UL25Δ44 Q72A** | 1:1 | 30/100 = 0.30 | 0.13 | 0.17 | 0.14 | 32/100 = 0.32 | 0.19 | 0.13 | 126.64 | 102.34 |
| | | 31/101 = 0.31 | 0.18 | 0.13 | | 27/99 = 0.27 | 0.09 | | 94.55 | |
| | | 31/105 = 0.30 | 0.18 | 0.12 | | 30/100 = 0.30 | 0.12 | | 85.84 | |
| | | 27/95 = 0.28 | 0.10 | 0.18 | 0.11 | 23/100 = 0.23 | 0.13 | 0.12 | 153.51 | 95.61 |
| | | 20/100 = 0.20 | 0.12 | 0.08 | | 25/100 = 0.25 | 0.13 | | 66.67 | |
| | | 23/100 = 0.23 | 0.15 | 0.08 | | 25/100 = 0.25 | 0.10 | | 66.67 | |
| | 1:6 | 23/100 =0.23 | 0.13 | 0.10 | 0.10 | 32/100 = 0.32 | 0.19 | 0.13 | 74.50 | 72.01 |
| | | 29/100 = 0.29 | 0.18 | 0.11 | | 27/99 = 0.27 | 0.09 | | 81.94 | |
| | | 26/100 = 0.26 | 0.18 | 0.08 | | 30/100 = 0.30 | 0.12 | | 59.59 | |
| | | 21/97 = 0.27 | 0.10 | 0.12 | 0.10 | 23/100 = 0.23 | 0.13 | 0.12 | 97.08 | 85.14 |
| | | 26/100 = 0.26 | 0.12 | 0.14 | | 25/100 = 0.25 | 0.13 | | 116.66 | |
| | | 20/100 = 0.20 | 0.15 | 0.05 | | 25/100 = 0.25 | 0.10 | | 41.66 | |
| | 1:8 | 23/99 = 0.23 | 0.17 | 0.06 | 0.08 | 32/96 = 0.33 | 0.16 | 0.16 | 37.47 | 47.53 |
| | | 24/98 = 0.24 | 0.17 | 0.07 | | 28/98 = 0.29 | 0.11 | | 45.02 | |
| | | 21/100 = 0.21 | 0.11 | 0.10 | | 33/100 = 0.33 | 0.22 | | 60.11 | |
| | | 25/95 = 0.26 | 0.13 | 0.13 | 0.09 | 37/100 = 0.37 | 0.24 | 0.20 | 65.77 | 43.33 |
| | | 29/100 = 0.29 | 0.17 | 0.12 | | 30/96 = 0.31 | 0.14 | | 59.27 | |
| | | 21/100 = 0.21 | 0.19 | 0.02 | | 39/94 = 0.41 | 0.22 | | 4.94 | |
| | 1:10 | 19/100 = 0.19 | 0.13 | 0.06 | 0.03 | 32/100 = 0.32 | 0.19 | 0.13 | 44.70 | 24.52 |
| | | 21/96 = 0.22 | 0.18 | 0.04 | | 27/99 = 0.27 | 0.09 | | 28.89 | |
| | | 18/105 = 0.18 | 0.18 | 0.00 | | 30/100 = 0.30 | 0.12 | | 0.00 | |
| | | 13/100 = 0.13 | 0.10 | 0.03 | 0.04 | 23/100 = 0.23 | 0.13 | 0.12 | 25.00 | 30.55 |
| | | 20/100 = 0.20 | 0.12 | 0.08 | | 25/100 = 0.25 | 0.13 | | 66.66 | |
| | | 15/100 = 0.15 | 0.15 | 0.00 | | 25/100 = 0.25 | 0.10 | | 0.00 | |
| | 1:20 | 23/99 = 0.23 | 0.17 | 0.06 | 0.05 | 32/96 = 0.33 | 0.16 | 0.16 | 37.47 | 28.51 |
| | | 19/100 = 0.19 | 0.17 | 0.02 | | 28/98 = 0.29 | 0.11 | | 12.02 | |
| | | 17/100 = 0.17 | 0.11 | 0.06 | | 33/100 = 0.33 | 0.22 | | 36.07 | |
| | | 23/100 = 0.23 | 0.13 | 0.10 | 0.37 | 37/100 = 0.37 | 0.24 | 0.20 | 49.39 | 18.11 |
| | | 17/100 = 0.17 | 0.17 | 0.00 | | 30/96 = 0.31 | 0.14 | | 0.00 | |

| | Ratio | ILVs/GUVs counted | Background (BG) | Raw Value (ILV-BG) | Average (AVG) | WT (ILVs/GUVs counted) | WT - BG | WT AVG | % Normalized Budding (Raw Value/WT AVG) * 100 | Biological Replicate Average (%) |
|---|---|---|---|---|---|---|---|---|---|---|
| | | 20/100 = 0.20 | 0.19 | 0.01 | | 39/94 = 0.41 | 0.22 | | 4.94 | |
| NEC-CBM220 + UL25Δ44 Q72A | CBM alone | 9/100 = 0.09 | 0.02 | 0.07 | | 11/100 = 0.11 | 0.09 | | 86.75 | |
| | | 10/100 = 0.10 | 0.01 | 0.09 | 0.08 | 7/98 = 0.07 | 0.06 | 0.08 | 111.53 | 103.54 |
| | | 10/100 = 0.10 | 0.01 | 0.09 | | 10/100 = 0.10 | 0.09 | | 112.34 | |
| | | 16/100 = 0.16 | 0.06 | 0.10 | | 19/103 = 0.18 | 0.12 | | 86.39 | |
| | | 24/100 = 0.24 | 0.11 | 0.13 | 0.12 | 21/102 = 0.21 | 0.10 | 0.11 | 112.31 | 105.66 |
| | | 24/100 = 0.24 | 0.10 | 0.14 | | 23/100 = 0.23 | 0.13 | | 118.28 | |
| | 1:1 | 9/100 = 0.9 | 0.02 | 0.07 | | 11/100 = 0.11 | 0.09 | | 86.74 | |
| | | 7/100 = 0.7 | 0.01 | 0.06 | 0.06 | 7/98 = 0.07 | 0.06 | 0.08 | 74.35 | 70.49 |
| | | 5/100 = 0.05 | 0.01 | 0.04 | | 10/100 = 0.10 | 0.09 | | 50.38 | |
| | | 25/100 = 0.25 | 0.06 | 0.19 | | 19/103 = 0.18 | 0.12 | | 164.14 | |
| | | 21/100 = 0.21 | 0.11 | 0.10 | 0.12 | 21/102 = 0.21 | 0.10 | 0.11 | 86.39 | 108.53 |
| | | 19/100 = 0.19 | 0.10 | 0.09 | | 23/100 = 0.23 | 0.13 | | 75.08 | |
| | 1:6 | 11/100 = 0.11 | 0.02 | 0.09 | | 11/100 = 0.11 | 0.09 | | 111.53 | |
| | | 10/100 = 0.10 | 0.01 | 0.09 | 0.07 | 7/98 = 0.07 | 0.06 | 0.08 | 111.53 | 93.19 |
| | | 5/91 = 0.05 | 0.01 | 0.04 | | 10/100 = 0.10 | 0.09 | | 56.50 | |
| | | 17/100 = 0.17 | 0.06 | 0.11 | | 19/103 = 0.18 | 0.12 | | 95.03 | |
| | | 21/100 = 0.21 | 0.11 | 0.10 | 0.10 | 21/102 = 0.21 | 0.10 | 0.11 | 86.39 | 88.38 |
| | | 20/100 = 0.20 | 0.10 | 0.10 | | 23/100 = 0.23 | 0.13 | | 83.72 | |
| | 1:10 | 5/100 = 0.05 | 0.02 | 0.03 | | 11/100 = 0.11 | 0.09 | | 37.17 | |
| | | 9/100 = 0.09 | 0.01 | 0.08 | 0.06 | 7/98 = 0.07 | 0.06 | 0.08 | 99.14 | 74.62 |
| | | 8/100 = 0.08 | 0.01 | 0.07 | | 10/100 = 0.10 | 0.09 | | 87.56 | |
| | | 16/98 = 0.16 | 0.06 | 0.10 | | 19/103 = 0.18 | 0.12 | | 89.21 | |
| | | 21/98 = 0.21 | 0.11 | 0.10 | 0.10 | 21/102 = 0.21 | 0.10 | 0.11 | 90.09 | 84.80 |
| | | 19/100 = 0.19 | 0.10 | 0.09 | | 23/100 = 0.23 | 0.13 | | 75.08 | |
| NEC220 + UL25Δ50 Q72A | 1:10 | 15/87 = 0.17 | 0.15 | 0.02 | | 34/98 =0.35 | 0.20 | | 10.33 | |
| | | 17/94 = 0.18 | 0.16 | 0.02 | 0.031 | 38/95 = 0.40 | 0.24 | 0.22 | 9.62 | 14.17 |
| | | 17/90 = 0.19 | 0.14 | 0.05 | | 35/99 = 0.35 | 0.21 | | 22.55 | |
| | | 12/100 = 0.12 | 0.05 | 0.07 | | 19/100 = 0.19 | 0.14 | | 43.75 | |
| | | 10/100 = 0.10 | 0.09 | 0.01 | 0.040 | 23/100 = 0.23 | 0.14 | 0.16 | 6.25 | 27.08 |
| | | 11/100 = 0.11 | 0.06 | 0.05 | | 26/100 = 0.26 | 0.20 | | 31.25 | |

| | Ratio | ILVs/GUVs counted | Background (BG) | Raw Value (ILV-BG) | Average (AVG) | WT (ILVs/GUVs counted) | WT - BG | WT AVG | % Normalized Budding (Raw Value/WT AVG) * 100 | Biological Replicate Average (%) |
|---|---|---|---|---|---|---|---|---|---|---|
| **NEC220 + UL25Δ58 Q72A** | 1:1 | 20/100 = 0.20 | 0.10 | 0.10 | 0.11 | 23/100 = 0.23 | 0.13 | 0.12 | 83.33 | 90.26 |
| | | 26/99 = 0.26 | 0.12 | 0.14 | | 25/100 = 0.25 | 0.13 | | 118.86 | |
| | | 23/99 = 0.23 | 0.15 | 0.08 | | 25/100 = 0.25 | 0.10 | | 68.60 | |
| | | 26/100 = 0.26 | 0.12 | 0.14 | 0.11 | 28/100 = 0.28 | 0.16 | 0.13 | 105.81 | 80.51 |
| | | 26/99 = 0.26 | 0.15 | 0.11 | | 28/100 = 0.28 | 0.13 | | 82.81 | |
| | | 24/100 = 0.24 | 0.17 | 0.07 | | 28/100 = 0.28 | 0.11 | | 52.90 | |
| | 1:6 | 19/100 = 0.19 | 0.10 | 0.090 | 0.12 | 23/100 = 0.23 | 0.13 | 0.12 | 75.00 | 96.30 |
| | | 24/90 = 0.27 | 0.12 | 0.15 | | 25/100 = 0.25 | 0.13 | | 122.22 | |
| | | 26/100 = 0.26 | 0.15 | 0.11 | | 25/100 = 0.25 | 0.10 | | 91.67 | |
| | | 28/96 = 0.29 | 0.12 | 0.17 | 0.10 | 28/100 = 0.28 | 0.16 | 0.13 | 129.74 | 76.82 |
| | | 21/95 = 0.22 | 0.15 | 0.07 | | 28/100 = 0.28 | 0.13 | | 51.39 | |
| | | 24/102 = 0.24 | 0.17 | 0.07 | | 28/100 = 0.28 | 0.11 | | 49.35 | |
| | 1:8 | 42/95 = 0.44 | 0.17 | 0.27 | 0.22 | 32/96 = 0.33 | 0.16 | 0.16 | 163.57 | 130.83 |
| | | 35/97 = 0.36 | 0.17 | 0.19 | | 28/98 = 0.29 | 0.11 | | 114.71 | |
| | | 30/100 = 0.30 | 0.11 | 0.19 | | 33/100 = 0.33 | 0.22 | | 114.21 | |
| | | 35/100 = 0.35 | 0.13 | 0.22 | 0.21 | 37/100 = 0.37 | 0.24 | 0.20 | 108.66 | 103.50 |
| | | 40/99 = 0.40 | 0.17 | 0.23 | | 30/96 = 0.31 | 0.14 | | 115.60 | |
| | | 35/96 = 0.36 | 0.19 | 0.17 | | 39/94 = 0.41 | 0.22 | | 86.23 | |
| | 1:10 | 17/99 = 0.17 | 0.10 | 0.072 | 0.085 | 23/100 = 0.23 | 0.13 | 0.12 | 59.76 | 70.80 |
| | | 19/100 = 0.19 | 0.12 | 0.070 | | 25/100 = 0.25 | 0.13 | | 58.33 | |
| | | 25/95 = 0.26 | 0.15 | 0.11 | | 25/100 = 0.25 | 0.10 | | 94.30 | |
| | | 28/100 = 0.28 | 0.12 | 0.16 | 0.093 | 28/100 = 0.28 | 0.16 | 0.13 | 120.93 | 69.77 |
| | | 22/100 = 0.22 | 0.15 | 0.07 | | 28/100 = 0.28 | 0.13 | | 50.59 | |
| | | 22/100 = 0.22 | 0.17 | 0.05 | | 28/100 = 0.28 | 0.11 | | 37.79 | |
| | 1:20 | 34/95 = 0.36 | 0.17 | 0.19 | 0.17 | 32/96 = 0.33 | 0.16 | 0.16 | 112.95 | 104.83 |
| | | 29/95 = 0.31 | 0.17 | 0.14 | | 28/98 = 0.29 | 0.11 | | 81.31 | |
| | | 31/100 = 0.31 | 0.11 | 0.20 | | 33/100 = 0.33 | 0.22 | | 120.23 | |
| | | 40/100 = 0.40 | 0.13 | 0.27 | 0.20 | 37/100 = 0.37 | 0.24 | 0.20 | 133.36 | 100.66 |
| | | 32/95 = 0.34 | 0.17 | 0.17 | | 30/96 = 0.31 | 0.14 | | 82.41 | |
| | | 35/96 = 0.36 | 0.19 | 0.17 | | 39/94 = 0.41 | 0.22 | | 86.23 | |

| | Ratio | ILVs/GUVs counted | Background (BG) | Raw Value (ILV-BG) | Average (AVG) | WT (ILVs/GUVs counted) | WT - BG | WT AVG | % Normalized Budding (Raw Value/WT AVG) * 100 | Biological Replicate Average (%) |
|---|---|---|---|---|---|---|---|---|---|---|
| **NEC220 + UL25Δ73** | 1:1 | 36/94 = 0.38 | 0.13 | 0.25 | 0.14 | 32/100 = 0.32 | 0.19 | 0.13 | 188.45 | 110.65 |
| | | 29/100 = 0.29 | 0.18 | 0.11 | | 27/99 = 0.27 | 0.09 | | 81.94 | |
| | | 26/99 = 0.26 | 0.18 | 0.08 | | 30/100 = 0.30 | 0.12 | | 61.55 | |
| | | 10/95 = 0.11 | 0.03 | 0.08 | 0.06 | 11/95 = 0.12 | 0.09 | 0.08 | 96.84 | 81.31 |
| | | 8/97 = 0.08 | 0.04 | 0.04 | | 9/97 = 0.09 | 0.05 | | 54.65 | |
| | | 9/98 = 0.09 | 0.02 | 0.07 | | 11/96 = 0.11 | 0.09 | | 92.43 | |
| | | 7/99 = 0.07 | 0.01 | 0.06 | 0.04 | 6/101 = 0.06 | 0.05 | 0.05 | 120.07 | 94.56 |
| | | 7/100 = 0.07 | 0.01 | 0.06 | | 7/105 = 0.07 | 0.06 | | 117.85 | |
| | | 6/98 = 0.06 | 0.04 | 0.02 | | 9/107 = 0.08 | 0.04 | | 45.75 | |
| | 1:6 | 27/100 = 0.27 | 0.13 | 0.14 | 0.96 | 32/100 = 0.32 | 0.19 | 0.13 | 104.29 | 72.93 |
| | | 31/100 = 0.31 | 0.18 | 0.13 | | 27/99 = 0.27 | 0.09 | | 96.84 | |
| | | 22/108 = 0.20 | 0.18 | 0.02 | | 30/100 = 0.30 | 0.12 | | 17.66 | |
| | | 12/100 = 0.12 | 0.03 | 0.09 | 0.07 | 11/95 = 0.12 | 0.09 | 0.08 | 115.80 | 95.15 |
| | | 10/100 = 0.10 | 0.04 | 0.06 | | 9/97 = 0.09 | 0.05 | | 77.20 | |
| | | 9/98 = 0.09 | 0.02 | 0.07 | | 11/96 = 0.11 | 0.09 | | 92.43 | |
| | | 7/100 = 0.07 | 0.01 | 0.06 | 0.05 | 6/101 = 0.06 | 0.05 | 0.05 | 118.70 | 101.68 |
| | | 6/101 = 0.06 | 0.01 | 0.05 | | 7/105 = 0.07 | 0.06 | | 96.90 | |
| | | 9/108 = 0.08 | 0.04 | 0.04 | | 9/107 = 0.08 | 0.04 | | 89.48 | |
| | 1:8 | 34/95 = 0.36 | 0.17 | 0.19 | 0.20 | 32/96 = 0.33 | 0.16 | 0.16 | 113.00 | 121.81 |
| | | 33/100 = 0.33 | 0.17 | 0.16 | | 28/98 = 0.29 | 0.11 | | 96.18 | |
| | | 37/100 = 0.37 | 0.11 | 0.26 | | 33/100 = 0.33 | 0.22 | | 156.30 | |
| | | 39/98 = 0.40 | 0.13 | 0.27 | 0.20 | 37/100 = 0.37 | 0.24 | 0.20 | 132.35 | 99.82 |
| | | 32/94 = 0.34 | 0.17 | 0.17 | | 30/96 = 0.31 | 0.14 | | 84.20 | |
| | | 34/95 = 0.36 | 0.19 | 0.17 | | 39/94 = 0.41 | 0.22 | | 82.93 | |
| | 1:10 | 35/99 = 0.35 | 0.13 | 0.22 | 0.14 | 32/100 = 0.32 | 0.19 | 0.13 | 166.52 | 105.02 |
| | | 26/95 = 0.27 | 0.18 | 0.09 | | 27/99 = 0.27 | 0.09 | | 69.80 | |
| | | 28/98 = 0.28 | 0.18 | 0.10 | | 30/100 = 0.30 | 0.12 | | 78.74 | |
| | | 11/93 = 0.12 | 0.03 | 0.09 | 0.66 | 11/95 = 0.12 | 0.09 | 0.08 | 113.59 | 86.24 |
| | | 8/100 = 0.08 | 0.04 | 0.04 | | 9/97 = 0.09 | 0.05 | | 51.47 | |
| | | 9/97 = 0.09 | 0.02 | 0.07 | | 11/96 = 0.11 | 0.09 | | 93.65 | |
| | | 5/113 = 0.04 | 0.01 | 0.03 | 0.05 | 6/101 = 0.06 | 0.05 | 0.05 | 67.74 | 92.97 |
| | | 9/119 = 0.08 | 0.01 | 0.07 | | 7/105 = 0.07 | 0.06 | | 129.00 | |

| | Ratio | ILVs/GUVs counted | Background (BG) | Raw Value (ILV-BG) | Average (AVG) | WT (ILVs/GUVs counted) | WT - BG | WT AVG | % Normalized Budding (Raw Value/WT AVG) * 100 | Biological Replicate Average (%) |
|---|---|---|---|---|---|---|---|---|---|---|
| | | 9/113 = 0.08 | 0.04 | 0.04 | | 9/107 = 0.08 | 0.04 | | 82.19 | |
| | 1:20 | 32/89 = 0.36 | 0.17 | 0.19 | 0.19 | 32/96 = 0.33 | 0.16 | 0.16 | 113.95 | 113.49 |
| | | 32/95 = 0.33 | 0.17 | 0.16 | | 28/98 = 0.29 | 0.11 | | 100.27 | |
| | | 32/100 = 0.32 | 0.11 | 0.21 | | 33/100 = 0.33 | 0.22 | | 126.24 | |
| | | 34/100 = 0.34 | 0.13 | 0.21 | 0.14 | 37/100 = 0.37 | 0.24 | 0.20 | 103.72 | 106.84 |
| | | 30/97 = 0.31 | 0.17 | 0.14 | | 30/96 = 0.31 | 0.14 | | 68.79 | |
| | | 27/100 = 0.27 | 0.19 | 0.08 | | 39/94 = 0.41 | 0.22 | | 39.51 | |
| | | 34/97 = 0.35 | 0.17 | 0.18 | | 30/96 = 0.31 | 0.14 | | 89.16 | |
| | | 37/98 = 0.38 | 0.19 | 0.19 | | 39/94 = 0.41 | 0.22 | | 92.63 | |
| **eGFP-UL25 Constructs** | | | | | | | | | | |
| **NEC220 + eGFP-UL25Δ50 Q72A** | 1:10 | 4/100 = 0.04 | 0.02 | 0.02 | 0.01 | 8/100 = 0.08 | 0.06 | 0.08 | 25.00 | 12.50 |
| | | 4/100 = 0.04 | 0.03 | 0.01 | | 15/100 = 0.15 | 0.12 | | 12.50 | |
| | | 3/100 = 0.03 | 0.03 | 0.00 | | 9/100 = 0.09 | 0.06 | | 0.00 | |
| | | 3/100 = 0.03 | 0.02 | 0.01 | 0.01 | 8/100 = 0.08 | 0.06 | 0.07 | 15.00 | 15.00 |
| | | 3/100 = 0.03 | 0.03 | 0.00 | | 9/100 = 0.09 | 0.06 | | 0.00 | |
| | | 0/100 = 0.00 | 0.02 | 0.02 | | 10/100 = 0.10 | 0.08 | | 30.00 | |
| **NEC-CBM220 + eGFP-UL25Δ50 Q72A** | 1:10 | 21/100 = 0.21 | 0.05 | 0.16 | 0.17 | 17/100 = 0.17 | 0.12 | 0.15 | 102.13 | 108.51 |
| | | 19/100 = 0.19 | 0.04 | 0.15 | | 21/100 = 0.21 | 0.17 | | 95.74 | |
| | | 22/100 = 0.22 | 0.02 | 0.20 | | 20/100 = 0.20 | 0.18 | | 127.66 | |
| | | 21/100 = 0.21 | 0.05 | 0.16 | 0.18 | 19/100 = 0.19 | 0.14 | 0.16 | 100.00 | 115.04 |
| | | 22/100 = 0.22 | 0.09 | 0.13 | | 23/100 = 0.23 | 0.14 | | 81.25 | |
| | | 29/90 = 0.32 | 0.06 | 0.26 | | 26/100 = 0.26 | 0.20 | | 163.88 | |

| | Ratio | ILVs/GUVs counted | Background (BG) | Raw Value (ILV-BG) | Average (AVG) | WT (ILVs/GUVs counted) | WT - BG | WT AVG | % Normalized Budding (Raw Value/WT AVG) * 100 | Biological Replicate Average (%) |
|---|---|---|---|---|---|---|---|---|---|---|
| NEC220 + eGFP-UL25Δ73 | 1:10 | 12/98 = 0.12 | 0.02 | 0.10 | 0.09 | 8/100 = 0.08 | 0.06 | 0.08 | 128.06 | 113.52 |
| | | 12/100 = 0.12 | 0.03 | 0.09 | | 15/100 = 0.15 | 0.12 | | 112.50 | |
| | | 11/100 = 0.11 | 0.03 | 0.08 | | 9/100 = 0.09 | 0.06 | | 100.00 | |
| | | 9/94 = 0.10 | 0.02 | 0.08 | 0.06 | 8/100 = 0.08 | 0.06 | 0.07 | 113.62 | 83.95 |
| | | 7/100 = 0.07 | 0.03 | 0.04 | | 9/100 = 0.09 | 0.06 | | 60.00 | |
| | | 7/97 = 0.07 | 0.02 | 0.05 | | 10/100 = 0.10 | 0.08 | | 78.25 | |
